# *N*-chlorination mediates protective and immunomodulatory effects of oxidized human plasma proteins

**DOI:** 10.1101/584961

**Authors:** Agnes Ulfig, Anton V. Schulz, Alexandra Müller, Natalie Lupilov, Lars I. Leichert

## Abstract

Hypochlorous acid (HOCl), a powerful antimicrobial oxidant, is produced by neutrophils to fight infections. Here we show that *N*-chlorination, induced by HOCl concentrations encountered at sites of inflammation, converts blood plasma proteins into chaperone-like holdases that protect other proteins from aggregation. This chaperone-like conversion was reversible by antioxidants and was abrogated by prior methylation of basic amino acids. Furthermore, reversible *N*-chlorination of basic amino acid side chains is the major factor that converts plasma proteins into efficient activators of immune cells. Finally, HOCl-modified serum albumin was found to act as a pro-survival molecule that protects neutrophils from cell death induced by highly immunogenic foreign antigens. We propose that activation and enhanced persistence of neutrophils mediated by HOCl-modified plasma proteins, resulting in the increased and prolonged generation of ROS, including HOCl, constitutes a potentially detrimental positive feedback loop that can only be attenuated through the reversible nature of the modification involved.

## INTRODUCTION

Recruitment and activation of neutrophils at sites of infection is considered one of the principal mechanisms by which the human body protects itself against diseases. The killing strategy of neutrophils involves the ingestion of pathogens into the phagosome, accompanied by the production of a diverse set of so-called reactive oxygen species (ROS), including superoxide anions (O_2_^.-^), hydrogen peroxide (H_2_O_2_) and hypochlorous acid (HOCl) in a process known as the respiratory burst (recently reviewed in refs. ^1,2^).

HOCl, a major inflammatory ROS, is produced from hydrogen peroxide and chloride ions by the heme enzyme myeloperoxidase (MPO) ^3,4^. The antimicrobial properties of HOCl are well documented and numerous reports have provided strong evidence for severe damage to bacterial components within the neutrophil phagosome ^5–7^.

A particularly important target of HOCl are proteins. In proteins, HOCl exposure typically leads to side chain modification ^8–12^, fragmentation ^13,14^, misfolding/aggregation ^15^ or intermolecular di-tyrosine cross-linking, a hallmark of HOCl-oxidized proteins ^16^. This often leads to their aggregation ^17–19^. Consequently, various mechanisms to counter the accumulation of these misfolded proteins under HOCl stress exist in bacteria. These are mediated by chaperones that are activated by HOCl, the very same reactive species they protect against. One of the first protective proteins to be noticed was Hsp33, which gets reversibly activated by HOCl through oxidation of four critical cysteine residues ^19^. This is not surprising, as cysteine thiols together with methionine residues react rapidly with HOCl ^20,21^. More recently, we found that *E. coli* RidA, a member of the highly conserved, but functionally diverse YjgF/YER057c/UK114 protein family, also undergoes HOCl-based conversion into a chaperone holdase. This chaperone was highly active as protector of proteins from HOCl-induced aggregation. However in this case, instead of cysteine oxidation, *N*-chlorination of basic amino acids was the mechanism of its activation ^17^, although reactivity of HOCl with side chains of lysine and arginine is four to seven orders of magnitude lower than that of cysteine^22^. But much like cysteine oxidation, *N*-chlorination is a reversible oxidative modification that can be removed by cellular antioxidants such as ascorbate, glutathione or thioredoxin and thus can switch the holdase function of RidA off ^17,23,24^.

Similar observations were made more recently with *E. coli* CnoX (YbbN) that, when activated via *N*-chlorination, binds to and prevents a variety of substrates from aggregation and being irreversible oxidized ^25^.

While HOCl is highly bactericidal, generation of HOCl by immune cells is not without risk to the human body itself (reviewed in refs. ^26–29^). During inflammatory processes, up to 30% of total cellular MPO is secreted by neutrophils into extracellular surroundings via degranulation, leakage during phagocytosis, or by association with NETs ^30^. Neutrophils, accumulated in the interstitial fluid of inflamed tissues, have been reported to generate HOCl at concentrations of up to 25-50 mM per hour ^31^. It is thus not surprising that HOCl can also drastically increase the activity of the extracellular chaperone α_2_-macroglobulin (α_2_M) in human blood plasma to counteract the aggregation of host proteins under hypochlorous acid stress ^32^. However, the underlying molecular mechanism of HOCl-mediated conversion of this plasma protein is still unclear.

And while neutrophils produce high amounts of HOCl, it does not accumulate at those levels, as it reacts instantly with diverse biological molecules including proteins, DNA ^33^, cholesterol ^34^ and lipids ^35^. But due to their high abundance in blood and interstitial fluid, human serum albumin (HSA) and other plasma proteins are considered the major targets of HOCl-mediated damage and as such constitute the main sink for HOCl in the vicinity of inflammation ^36–40^. The resulting products of the reaction of plasma proteins with hypochlorous acid, known as advanced oxidation protein products (AOPPs), have, therefore, been employed as *in vivo* markers of chronic inflammation ^41^. Accumulation of AOPPs has been first discovered in patients with chronic kidney disease ^41^ and later also found in a variety of other inflammatory diseases, e.g. cardiovascular disease, neurodegenerative disorders, rheumatoid arthritis and some cancers (recently reviewed in ref. ^42^).

To date, a number of studies have been carried out to elucidate the role of AOPPs in inflammatory processes ^43–46^. Accumulating experimental evidence supports a critical contribution of AOPPs to the progression of inflammation ^44^. HOCl-modifed HSA accumulates in inflammatory diseases and was found to act as proinflammatory mediator by increasing oxidative stress and inflammation through stimulation of leukocytes ^46,47^.

Based on these findings we hypothesized that reversible *N*-chlorination in response to HOCl-stress could be the principal chemical modification contributing to the observed physiological properties of AOPPs. Furthermore, as we observed in previous studies that the HSA homologue bovine serum albumin (BSA) could be transformed into a chaperone-like holdase by HOCl-treatment ^17^, we speculated that *N*-chlorination serves as general rather than specific mechanism to transform certain proteins into a chaperone-like state.

Here, we study the effects of reversible *N*-chlorination on the function of human plasma proteins. We show that, upon *N*-chlorination, not only α_2_M, but all plasma fractions tested, exhibit chaperone-like activity and as such could prevent the HOCl-induced formation of protein aggregates at the site of inflammation. Moreover, exposure to HOCl at concentrations present in chronically inflamed tissues turned the majority of plasma proteins into efficient activators of neutrophil-like cells. Previous studies revealed that HOCl-treated HSA can stimulate leukocytes to produce more ROS during inflammation ^47,48^. Now we find that reversible *N*-chlorination is the main chemical modification that mediates the activation of NADPH oxidase-dependent ROS generation by immune cells by AOPPs. Finally, we show that HOCl-modified HSA is a pro-survival factor for immune cells and protects neutrophils from cell death by highly immunogenic antigens.

Our data strongly suggest that *in vivo* reversible *N*-chlorination of human plasma proteins not only converts them into effective chaperone-like holdases but is also the principal mechanism that turns these proteins into mediators of the innate immune system.

## MATERIAL AND METHODS

### Preparation of plasma protein solutions

Albumin from human serum (HSA, Product # A9511), human γ-globulins (Product # G4386), immunoglobulin G from human serum (IgG, Product # I4506) Cohn fraction IV (Product # G3637) and human whole serum (sterile-filtered from male AB plasma, Product # H4522) were purchased from Sigma-Aldrich, St. Louis, USA, and used without further purification.

Protein stock solutions were freshly prepared by dissolving or diluting varying amounts of the protein fractions in 1×PBS buffer, pH 7.4 (Gibco Life Sciences).

### Purification of α_2_-macroglobulin

α_2_-Macroglobulin was purified from human plasma (obtained from Zen-Bio, Inc., North Carolina, USA, Product # SER-SLP, Lot # 11108) by the method of Imber and Pizzo, 1981 ^49^ with slight modifications. Briefly, 880 mL fresh-frozen human plasma were thawed on ice and dialyzed against frequent changes of deionized water for 72 hours at 4 °C using a Spectra/Por dialysis membrane with a MWCO of 12-14 kDa (Spectrum Laboratories Inc., Rancho Dominguez, CA). Insoluble material was removed by centrifugation for 30 minutes at 10,000 × g and 4 °C. The supernatant plasma solution (200 mL) was then dialyzed for 24 hours at 4 °C against 5 L 1×PBS pH 6.0. Metal chelate chromatography was performed at 4 °C using an IMAC zinc-Sepharose 6 Fast Flow column (2.6 × 20 cm, GE Healthcare Life Sciences, Amersham, UK) equilibrated with 1×PBS pH 6.0. Dialyzed plasma was applied to the column and washed with 1×PBS pH 6.0 until the absorption at 280 nm of the eluant (measured with a JASCO V-650 UV/VIS spectrophotometer (JASCO, Tokyo, Japan)) reached a value lower than 0.01. Bound protein was then eluted from the column with 0.01 M NaOAc, 0.15 M NaCl, pH 5.0. Peak protein fractions were combined and concentrated using Vivaspin 20 concentrators with a MWCO of 100 kDa. Gel filtration of the concentrated protein pool fraction was performed at 4 °C on a HiPrep 26/60 Sephacryl S-300 High resolution column (2.6 × 60 cm) equilibrated with 1×PBS pH 7.4. High molecular weight peak fractions containing α_2_-macroglobulin were combined, concentrated and dialyzed against 1×PBS pH 7.4 containing 40% (v/v) glycerol. Aliquots were stored at −20° C.

### Determination of protein concentrations

Protein concentration in g ⋅ L^−1^ of human serum was calculated using Pierce^TM^ Bicinchoninic Acid (BCA) Protein Assay Kit (Thermo Fisher Scientific, Waltham, Massachusetts, USA) with bovine albumin as standard carried out following the manufacturer’s instructions. To calculate a molar concentration, an average molar mass of proteins of 66,357.12 Da was assumed (molar mass of human serum albumin).

For HSA, IgG, the γ-globulin fraction, and α_2_-macroglobulin, the concentration was determined by measuring the absorbance at 280 nm (A_280nm_) using a JASCO V-650 UV/VIS spectrophotometer. The molar extinction coefficient used for HSA was ε_280_= 35,700 M^−1^ cm^−1^ ^50^. Concentration of the γ-globulin fraction was estimated using the extinction coefficient for immunoglobulin G (IgG) of 1.36 cm^−1^ (mg ⋅ mL^−1^)^−1^ ^51^. Assuming a molecular weight of 150,000 Da ^52^, the molar extinction coefficient at 280 nm used for IgG was 210,000 M^−1^ cm^−1^. The molar extinction coefficient used for α_2_-macroglobulin, ε_280_= 145,440 M^-1^ cm^-1^, was calculated from amino acid sequence using ProtParam ^53^.

Concentration of Cohn fraction IV was determined using Pierce^TM^ Bicinchoninic Acid (BCA) Protein Assay Kit and bovine serum albumin (BSA) as standard according to the manufacturer’s instructions. To calculate a molar concentration, a weighted average protein mass of 80,000 Da was assumed based on the composition of the Cohn fraction IV.

### Methylation of proteins

Proteins were dissolved in 1ml 1×PBS pH 7.4 to a concentration of 10 mg ⋅ mL^−1^ and the solution cooled to 4 °C. 20 µl of 60 mg ⋅ mL^−1^ dimethylamine borane complex and 40 µl 1M formaldehyde were then added. After 2 hours of incubation at 4 °C, this step was repeated. 2 hours later a final aliquot of 10 µl dimethylamine borane complex solution was added, before incubation of the reaction mixture at 4 °C overnight. The next morning 125 µl of 1M Tris pH 7.5 were added to stop the reaction. The reacting agents were then separated from the now methylated proteins by size-exclusion chromatography using Nap^TM^-5 columns according to the manufacturer’s instructions (GE Healthcare Life Sciences, Amersham, UK).

### HOCl-treatment of proteins and reduction

The concentration of the NaOCl stock solution of 0.64 M (Sigma-Aldrich, St. Louis, USA) was confirmed regularly by measuring the absorbance at 292 nm using a JASCO V-650 UV/VIS spectrophotometer and the extinction coefficient ε_292_= 350 M^−1^ cm^−1^. When necessary, NaOCl stock solution was diluted by mixing an adequate volume of NaOCl with 1× PBS solution pH 7.4 immediately prior to each chlorination reaction.

Varying amounts of a protein were then treated with a 10-fold, 50-fold and/or 150-fold molar excess of NaOCl for 10 minutes at 30 °C (maximum NaOCl concentration used was 50 mM – a concentration that can be produced by neutrophils per hour in chronically inflamed tissues ^31^). Excess HOCl was removed by size-exclusion chromatography using Nap^TM^-5 columns according to the manufacturer’s instructions. Due to dilution during the Nap^TM^-5 desalting step, protein concentrations were re-determined as described above.

To reverse protein *N*-chlorination in HOCl-treated proteins, sodium ascorbate was dissolved in 1×PBS pH 7.4 to a concentration of 1 M and the proteins were incubated with a 50-fold molar excess of sodium ascorbate for 45 minutes at 37 °C. After removal of excess reductant (see above), protein concentrations were again re-determined.

### Protein aggregation assays with citrate synthase

200 µL Citrate synthase in ammonium sulfate solution (Sigma-Aldrich, St. Louis, USA) was dialyzed overnight against 2 L 20mM Tris 2mM EDTA buffer at 4°C under constant stirring using a Spectra/Por dialysis membrane with a MWCO of 6-8,000 Da. (Spectrum Laboratories Inc., Rancho Dominguez, CA). This dialyzed citrate synthase preparation was then chemically denatured in 4.5 M GdnHCl, 40 mM HEPES, pH 7.5 at room temperature overnight. The final concentration of denatured citrate synthase was 12 μM. Aggregation was induced by the addition of 20 µL denatured citrate synthase stock to 1580 μL 40 mM HEPES/KOH buffer, pH 7.5. Final concentration of citrate synthase in the aggregation assay was thus 0.15 μM. Untreated or treated plasma proteins were added to the assay buffer to a final concentration of 0.5 μM (α_2_-Macroglobulin) or 1.5 μM (HSA, IgG, γ-globulins, Cohn fraction IV) (corresponding to a 3.3-fold and 10-fold molar excess over the dimeric citrate synthase, respectively) prior to the addition of citrate synthase. The increase of light scattering was monitored in a JASCO FP-8500 fluorescence spectrometer equipped with an EHC-813 temperature-controlled sample holder (JASCO, Tokyo, Japan) at 30 °C for 200-240 s under continuous stirring. Measurement parameters were set to 360 nm (Ex/Em), 2.5 nm slit width (Ex/Em) and medium sensitivity. Chaperone activity was expressed as the difference between initial and final light scattering of an individual sample in arbitrary units. Aggregation of citrate synthase in the absence of any other proteins was set to 100%. Depending on the batch of citrate synthase, absolute maximum and minimum light scattering values may vary, thus control experiments with the same batch were carried out for each individual aggregation experiment.

### Detection of accessible amino groups in proteins

Accessible amino groups in HSA were detected using fluorescamine (Sigma-Aldrich, St. Louis, USA) as described ^54^. Briefly, 334 μl of 3 mg ⋅ mL^−1^ fluorescamine stock in acetone were added to 1 mL of 80 μg ⋅ mL^−1^ native HSA or the variously treated HSA solutions described above. Emission spectrum of fluorescamine from 400 to 600 nm was measured upon excitation with 388 nm using a JASCO FP-8500 fluorescence spectrometer. The relative amount of accessible amino groups in the variously treated HSA samples was calculated by setting the maximum fluorescence of native HSA to 100% representing the total relative amino group content.

### Nile red hydrophobicity assay

Nile red (Sigma-Aldrich, St. Louis, USA) was dissolved in dimethyl sulfoxide (DMSO) to a final concentration of 30 μM. Varying amounts of native or HOCl-treated HSA (0-200 μM) in 1×PBS were mixed with Nile red stock to a final dye concentration of 1.6 μM. Fluorescence was measured using a JASCO FP-8500 fluorescence spectrometer with the following parameters: 550 nm excitation, 570-700 nm emission, 5 nm slit width (Ex/Em) and medium sensitivity. Concentrations of native or HOCl-treated HSA, at which the proteins were half-saturated with dye were calculated by plotting the fluorescence intensity of Nile red against the logarithm of the molar protein concentrations. Data were fitted with GraphPad Prism 8 software using the sigmoid fit function.

### PLB-985 culture and differentiation

The human myeloid leukemia cell line PLB-985 (certified mycoplasma negative, obtained from DSMZ, German collection of microorganisms and cell culture) was cultured in RPMI-1640 medium supplemented with 10% heat-inactivated FBS and 1% GlutaMAX (Life Technologies, Carlsbad, CA) at 37°C with 5% CO_2_. Cells were passaged twice weekly to maintain a cell density between 2 × 10^5^ and 1 × 10^6^ ⋅ mL^−1^ and used until passage no. 10. For differentiation into neutrophil-like cells, cells were seeded at a density of 2 × 10^5^ ⋅ mL^−1^ and cultured for 96 hours in RPMI-1640 medium with 10% FBS, 1% GlutaMAX and 1.25% DMSO ^55^. After 72 hours of incubation in the presence of DMSO, 2000 U ⋅ mL^−1^ interferon-γ (ImmunoTools, Friesoythe, Germany) was added to the cell culture ^56^. In a previous study, differentiation was checked by detecting the expression of the associated surface markers CD11b and CD64 (see ref. ^7^). The viability of the cells was evaluated using trypan blue dye and was typically >90%.

### Chemiluminescence-based NADPH-oxidase activity assay

NADPH-oxidase-dependent superoxide production was selectively measured by chemiluminescence (CL) using the chemiluminogenic substrate lucigenin (10,10’-Dimethyl-9,9’-biacridinium dinitrate; Carl Roth, Karlsruhe, Germany) ^57^.

Differentiated PLB-985 cells were washed once with 1×PBS pH 7.4 and diluted in the same buffer to a final concentration of 5 × 10^6^ cells ⋅ mL^−1^. 100 µL of this cell suspension were placed in the wells of a non-transparent, white, clear-bottom 96-well plate (Nunc, Rochester, NY). For some experiments, cells were preincubated with 10 µM diphenyleneiodium (DPI; NADPH oxidase inhibitor), 100 nM wortmannin (phosphoinositide 3-kinase (PI3K) inhibitor), 200 nM Gö 6983 (protein kinase C (PKC) inhibitor) or vehicle (1% DMSO) for 30 minutes at 37 °C. All inhibitors dissolved in DMSO were purchased from Sigma-Aldrich, St. Louis, USA. The final concentration of DMSO in all wells was adjusted to 1%. The bottom of the plate was covered using a white, non-transparent adhesive seal prior to measurement. 50 µL of either 1×PBS (resting CL) or agents to be tested, including the untreated and treated plasma proteins were added to the respective wells. Final concentrations were: 2 mg ⋅ mL^−1^ α_2_-macroglobulin, 3 mg ⋅ mL^−1^ HSA, IgG, γ-globulins and Cohn fraction IV, 0.2 µM PMA (Phorbol 12-myristate 13-acetate; Sigma-Aldrich, St. Louis, USA). Lucigenin was dissolved in 1×PBS to a concentration of 400 µM immediately prior to measurement and 50 µL were added to the wells. Chemiluminescence was measured every 1-2 minutes over 1.5 hours at 37 °C using the Synergy H1 multi-detection microplate reader (Biotek, Bad Friedrichshall, Germany) in triplicates and chemiluminescence activity was expressed as integrated total counts as calculated by the addition of rectangles with unit width under individual data points. In inhibition assays, NADPH-oxidase activation, as measured by chemiluminescence activity, induced by PMA, HSA, treated with a 50-fold molar excess of HOCl, or IgG, treated with a 150-fold molar excess of HOCl, was set to 100% and other relevant data as percentage of this control.

### Cloning of Ag85B

*E. coli* strains, plasmids and primers used in this study are listed in Table 1. Ag85B gene (*fbpB*) of *Mycobacterium bovis* (differs by one base-pair from corresponding gene of *M. tuberculosis*, resulting in a Leu to Phe replacement at position 100 ^58^) was designed with optimized codon usage for expression in *E. coli*, synthesized and cloned into pEX-A2 standard vector by Eurofins Genomics. *fbpB* was amplified from pEXA_fbpB by PCR using primers fbpB-fw and fbpB-rv, purified according to the instructions of the NucleoSpin Gel and PCR Clean-up Kit (Macherey-Nagel GmbH, Düren, Germany) and cloned into the pET22b(+) expression vector via the restriction sites NdeI and XhoI with a hexahistidine (His_6_)-tag placed at the C-terminal end of the gene. *E. coli* DH5α cells were transformed with the plasmid using a standard heat-shock method and plated on Luria Bertani (LB) agar plates supplemented with 50 µg ⋅ mL^−1^ ampicillin. Screening for recombinant plasmids was performed by colony PCR, followed by isolation of potentially correct plasmids from the respective strains. Successful cloning of *fbpB* gene into pET22b(+) vector was verified by sequencing.

**Table 1:**
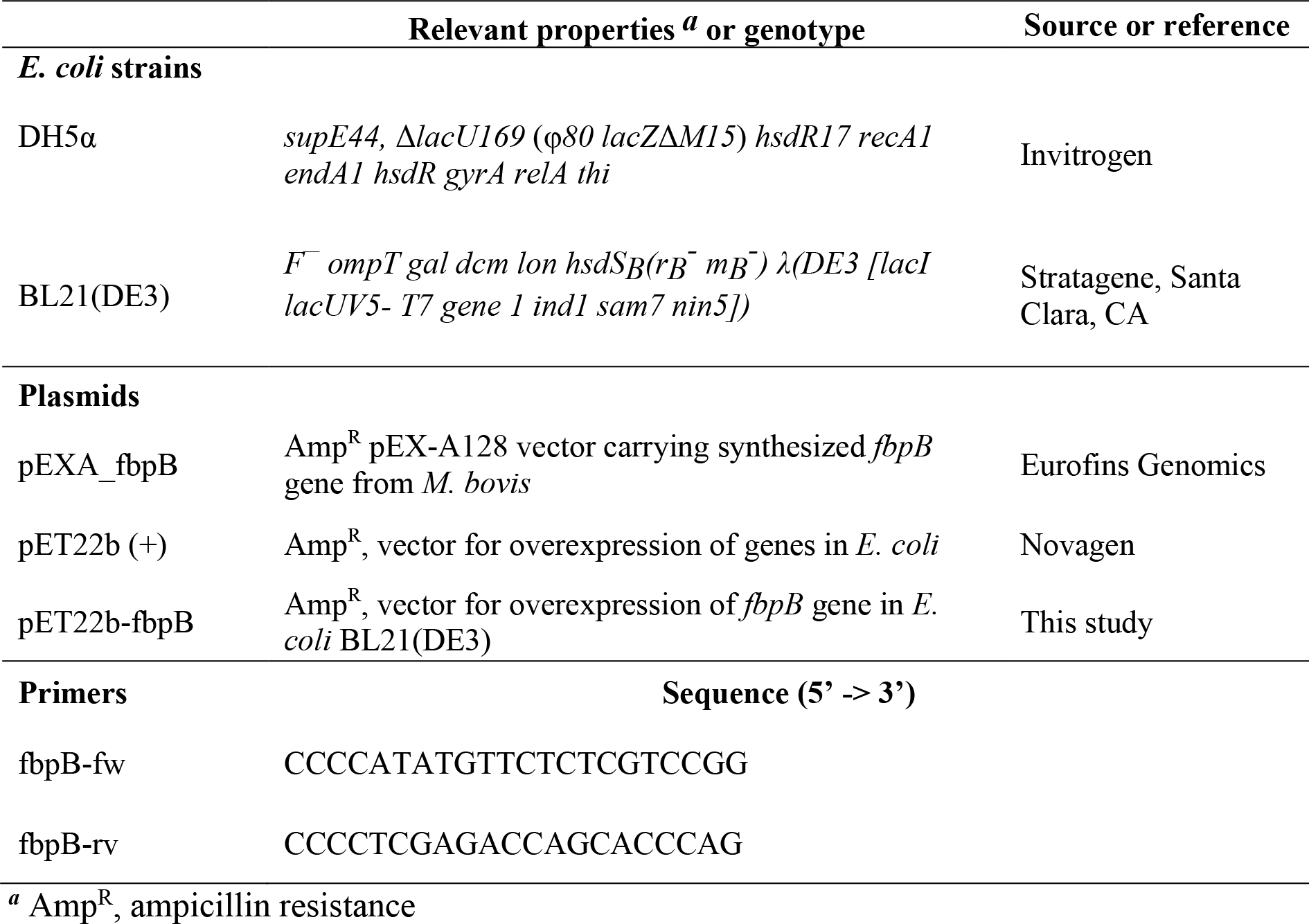
*E. coli* strains, plasmids and primers used in this study.

### Expression and purification of Ag85B

For heterologous expression and subsequent purification of His_6_-tagged Ag85B, recombinant plasmid pET22b-fbpB was transferred into the *E. coli* expression strain BL21 (DE3) (see Table 1). The transformed cells were plated on LB agar plates containing 50 µg ⋅ mL^−1^ ampicillin and incubated at 37 °C for 24 hours. 2 × 50 ml LB medium with ampicillin were inoculated with a single colony from the agar plate and incubated at 37 °C overnight. These overnight cultures were then used for inoculation of 5 × 1 L ampicillin-containing LB medium to a starting optical density at 600 nm (OD_600_) of 0.1. The bacteria were grown at 37 °C with shaking at 130 rpm until the OD_600_ was ~ 0.5. Expression of Ag85B including a C-terminal hexahistidine (His_6_)-tag was induced with 1 mM isopropyl-beta-D-thiogalactopyranoside (IPTG) and continued for ~ 12 hours (overnight) at 20 °C. Cells were harvested by centrifugation at 6500 × g for 45 minutes at 4 °C. Pellets were washed once with 1×PBS and resuspended in lysis buffer (5 mM imidazole, 300 mM NaCl, 50 mM NaH_2_PO_4_, pH 8.0). Cells were disrupted using a Constant Cell disrupter (Constant Systems Limited, Daventry, England), and the obtained lysate was centrifuged for 45 minutes at 4 °C and 57,500 × g. Solid precipitate and the supernatant were separated and evaluated for detection of recombinant protein by SDS-PAGE, followed by Coomassie Blue G-250 staining. Overexpressed Ag85B protein was found predominantly in the pellet fraction. Pellet was thus resuspended in lysis buffer containing 8 M urea and thoroughly homogenized. Suspension was applied to a polystyrene column filled with nickel-nitrilotriacetic (Ni-NTA) resin, and washed with urea-containing lysis buffer, followed by a washing step with wash buffer supplemented with a small amount of imidazole (8M urea, 20 mM imidazole, 300 mM NaCl, 50 mM NaH_2_PO_4_, pH 8.0). Recombinant Ag85B was eluted from the column using 250 mM imidazole buffer solution (8M urea, 250 mM imidazole, 300 mM NaCl, 50 mM NaH_2_PO_4_, pH 8.0). In order to remove imidazole, combined elution fraction was diluted 1:10 in sodium phosphate buffer (300 mM NaCl, 50 mM NaH_2_PO_4_, pH 8.0) and mixed overnight at 4 °C. Precipitated protein was collected by centrifugation for 45 minutes at 4 °C and 57,500 × g and resuspended in urea-containing sodium phosphate buffer (8M urea, 300 mM NaCl, 50 mM NaH_2_PO_4_, pH 8.0). Concentration of purified Ag85B protein was quantified spectrophotometrically by measuring the absorbance at 280 nm using a JASCO V-650 UV/VIS spectrophotometer. The molar extinction coefficient used for Ag85B was ε_280_= 75,860 M^−1^ cm^−1^ ^59^.

### Flow cytometry-based cell apoptosis assay

Differentiated PLB-985 cells were counted, washed once with 1×PBS pH 7.4 and diluted in RPMI-1640 medium supplemented with 10% heat-inactivated FBS to a final concentration of 3.3 × 10^6^ cells ⋅ mL^−1^. 600 µL of this cell suspension were placed in the wells of a transparent, flat-bottom 24-well plate (Sarstedt, Nümbrecht, Germany). The cells were preincubated with 100 nM wortmannin, 50 µM Z-VAD-FMK, a broad spectrum pan-caspase inhibitor (dissolved in DMSO; Santa Cruz Biotechnology, Dallas, TX, USA), 250 µM cytochalasin D, an inhibitor of actin polymerization (dissolved in DMSO; Sigma-Aldrich, St. Louis, USA), 155 µM native or modified HSA, that has been treated with a 50-fold molar excess of HOCl as described above, or vehicle (1% DMSO) for 1 hour at 37 °C. The final concentration of DMSO in all wells was adjusted to 1%. Subsequently, 1 µM Ag85B (85 µM stock in urea-containing buffer (8 M urea, 300 mM NaCl, 50 mM NaH_2_PO_4_, pH 8.0)), 2 µM staurosporine (dissolved in DMSO; Sigma-Aldrich, St. Louis, USA) or vehicle (94 mM urea and 1% DMSO) were added. After 1 hour of incubation with Ag85B or 6 hours with staurosporine at 37°C, cells were washed once with cold 1×PBS, followed by one washing step with 1× Annexin V binding buffer. For the analysis of cell viability, the variously treated cell suspensions were stained with Annexin V-FITC and propidium iodide (PI) using the Dead Cell Apoptosis Kit (Invitrogen, Thermo Fisher Scientific, Waltham, Massachusetts, USA) according to the manufacturer’s instructions and subsequently subjected to flow cytometry. Samples were analyzed using a BD FACSCanto II flow cytometer (Becton, Dickinson and Company). Fluorescence emitted by Annexin V-FITC and PI was measured through a 530/30-nm and 585/42-nm bandpass filter, respectively, upon excitation with an argon ion laser operating at 488 nm. Single-stained compensation controls were used to calculate the compensation matrix. 20,000 events were acquired and recorded per sample. Data were analyzed using FlowJo (version 10) software.

### Fluorescent labeling of Ag85B

For fluorescent labeling of Ag85B the green fluorescent dye CF^TM^ 488A succinimidyl ester (Sigma-Aldrich, St. Louis, USA) was used. Labeling occurred via reaction of the succinimidyl ester group of the dye with amine groups of Ag85B. Labeling was performed according to the manufacturer’s instructions with slight modifications. To prevent aggregation of Ag85B, labeling reaction was performed in urea-containing buffer (8 M urea, 300 mM NaCl, 50 mM NaH_2_PO_4_, pH 8.0) and carried out for 6 hours at room temperature under continuous shaking. Excess dye was removed by size-exclusion chromatography using PD-10 desalting columns containing Sephadex G-25 resin according to the manufacturer’s instructions (GE Healthcare Life Sciences, Amersham, UK). Concentration of the conjugate and degree of labeling (DOL) were calculated using the formula provided by the dye’s manufacturer. DOL was 3.1.

### Ag85B-488 uptake assay

For the antigen uptake assay, differentiated PLB-985 cells were incubated with fluorescently-labeled Ag85B (Ag85B-488) protein in the presence or absence of native and HOCl-modified HSA in the same way as for the cell apoptosis assay described above. After 1 hour of incubation at 37°C, cells were washed twice with 1×PBS, followed by fixation with 4% paraformaldehyde (PFA) for 10 minutes on ice. Cells were washed with 1×PBS and subsequently subjected to flow cytometry. Samples were analyzed using a BD FACSCanto II flow cytometer. Fluorescence emitted by Ag85B-488 was measured through a 530/30-nm bandpass filter upon excitation with the 488 nm argon laser line. 30,000 events were acquired and recorded per sample. Data were analyzed using FlowJo (version 10) software.

### Ag85B aggregation assay

Aggregation of Ag85B was induced by the stepwise addition of 75 µL Ag85B stock (5 × 15 µL every 20 seconds) to 1525 μl 40 mM HEPES/KOH buffer, pH 7.5. Final concentration of Ag85B in the aggregation assay was 0.186 μM. Native or variously treated HSA as described above, was added to the assay buffer to a final concentration of 14.88 μM (corresponding to a 80-fold molar excess over Ag85B) prior to the addition of Ag85B. The increase of light scattering was monitored in a JASCO FP-8500 fluorescence spectrometer equipped with an EHC-813 temperature-controlled sample holder at 30 °C for 200 s under continuous stirring. Measurement parameters were set to 360 nm (Ex/Em), 2.5 nm slit width (Ex/Em) and medium sensitivity. Chaperone activity was expressed as the difference between initial and final light scattering of an individual sample in arbitrary units. Aggregation of Ag85B in the absence of HSA was set to 100%.

## RESULTS

### HOCl-treated human serum prevents protein aggregation

In previous experiments we showed that the bacterial protein RidA is transformed into a competent holdase-type chaperone upon treatment with HOCl or *N*-chloramine ^17^. In control experiments, we discovered that bovine serum albumin also shows increased chaperone activity in response to HOCl ^17^. Since HOCl can be present in a physio-pathological context, most notably in the vicinity of inflammation, we wanted to test if human serum proteins can also be transformed into holdase-type chaperones through HOCl-treatment. Thus, we performed aggregation assays using chemically denatured citrate synthase and untreated and HOCl-treated human serum. Human serum was incubated with an estimated 10-fold molar excess of HOCl for 10 minutes at 30°C, a concentration that was sufficient to fully activate or markedly improve the chaperone function of RidA and BSA, respectively ^17^. When chemically denatured citrate synthase was diluted into denaturant-free buffer, it formed aggregates that can be monitored by increased light scattering of the solution (Fig. 1). This aggregation of citrate synthase was not prevented by the addition of untreated human serum. However, when pre-incubated with a 10-fold molar excess of HOCl, serum significantly decreased aggregate formation. This suggested to us, that at least some serum proteins could be transformed to a holdase-type chaperone by *N*-chlorination in a mechanism similar to RidA. Because *N*-chlorination can be reduced by certain antioxidants, we used ascorbate to re-reduce HOCl-treated serum. Ascorbate is a mild antioxidant, which typically does not reduce native or HOCl-induced disulfide bonds. This ascorbate-treated serum lost its capability to bind denatured citrate synthase, suggesting an *N-*chlorination-based mechanism.

**Figure 1:**
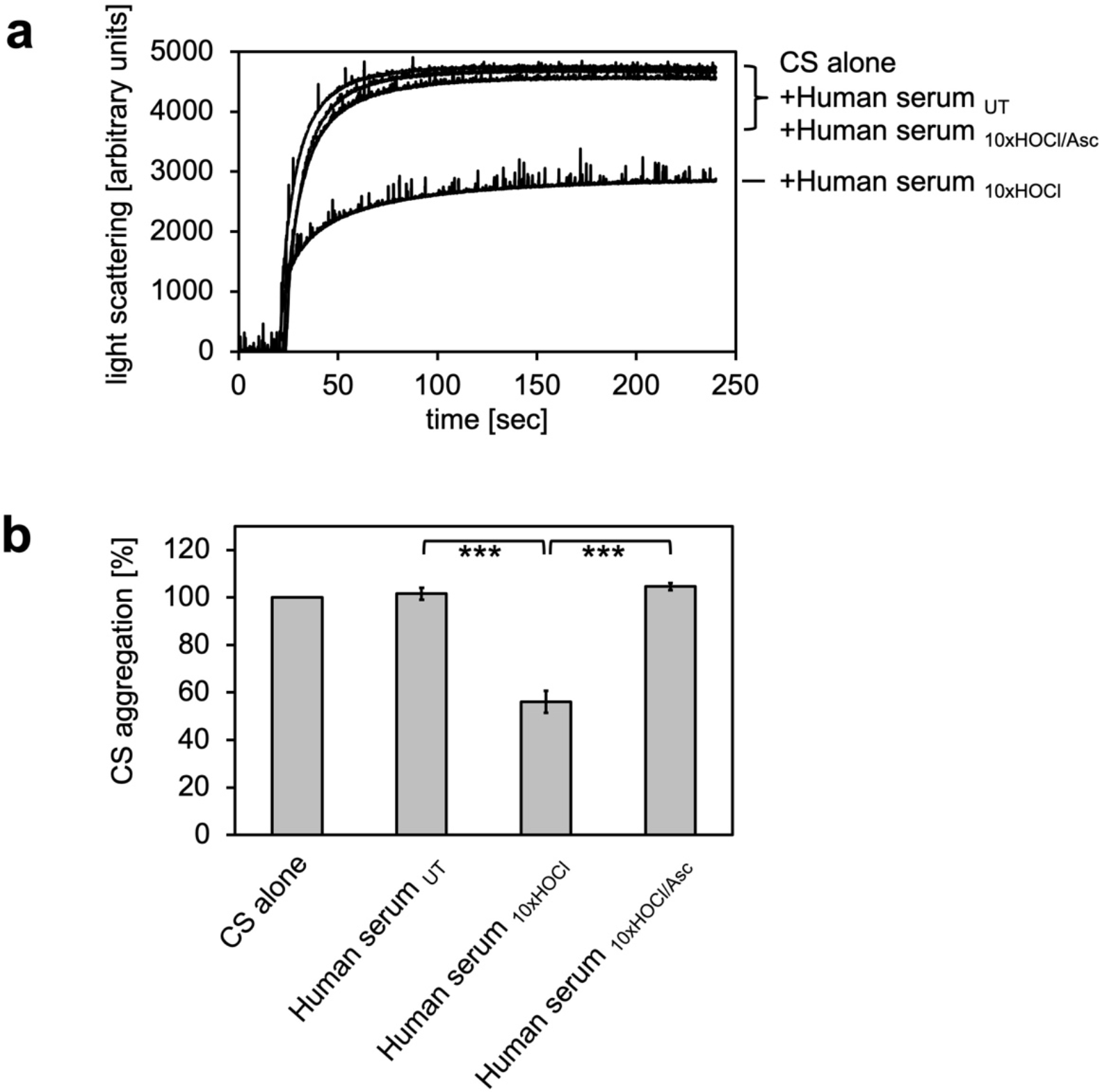
HOCl-treated human serum decreases protein aggregation. Human serum, when treated with a 10-fold molar excess of HOCl (Human serum _10×HOCl_), significantly decreases aggregation of chemically-denatured citrate synthase as measured by light scattering at 360 nm. Reduction of HOCl-treated human serum with a 50-fold molar excess of the antioxidant ascorbate (Human serum _10×HOCl/Asc_) reverses this chaperone-like conversion of the serum. **(a)** A representative measurement of citrate synthase aggregation in the presence of untreated (Human serum _UT_), HOCl-treated (Human serum _10×HOCl_) and re-reduced (Human serum _10×HOCl/Asc_) human serum is shown. **(b)** Data are represented as means and standard deviations from three independent aggregation assays. Student’s t-test: ***p < 0.001. Aggregation of citrate synthase in the absence of human serum was set to 100% and all the data are presented as percentage of this control. Labels of aggregation curves are written in the order of the final intensity of light scattering of the respective treatment.

### Albumin, the major protein component of serum, shows HOCl-induced chaperone-like activity

The major protein component of human serum is human serum albumin (HSA). Previously, we showed that its bovine homologue bovine serum albumin (BSA) exhibits increased chaperone activity upon treatment with HOCl ^17^. We therefore suspected that HSA could be a major contributor to the observed decrease in protein aggregate formation in the presence of HOCl-treated serum. To test whether HSA gains chaperone function upon exposure to HOCl, HSA at a final concentration of 1 mM was treated with a 10-fold molar excess of HOCl for 10 minutes at 30 °C. An HOCl concentration of at least 10 mM has been shown to be required for the generation of significant quantities of the so-called advanced oxidation protein products (AOPPs) in plasma, associated with a number of inflammatory diseases ^41^. The localized concentration of HOCl generated by accumulated neutrophils in the interstitium of chronically inflamed tissues, however, can be much higher and reach values of 25-50 mM per hour ^31^. To mimic a state of chronic inflammation, HSA was thus also incubated in the presence of a 50-fold molar excess of HOCl corresponding to the maximum reported concentration of 50 mM. HSA treated in both ways significantly reduced the aggregation of citrate synthase when added at a 10-fold molar excess, showing that HSA acts as a potent chaperone in serum upon modification by HOCl (Fig. 2 a, b). The chaperone activity of HSA, however, was much higher upon exposure to a 50-fold molar excess of HOCl, suggesting a dose-dependent activation of the HSA chaperone function by HOCl. In contrast to bacterial RidA and Hsp33 ^17,19^, full activation of the chaperone function of HSA seems to require a higher oxidation/chlorination level of the protein.

**Figure 2:**
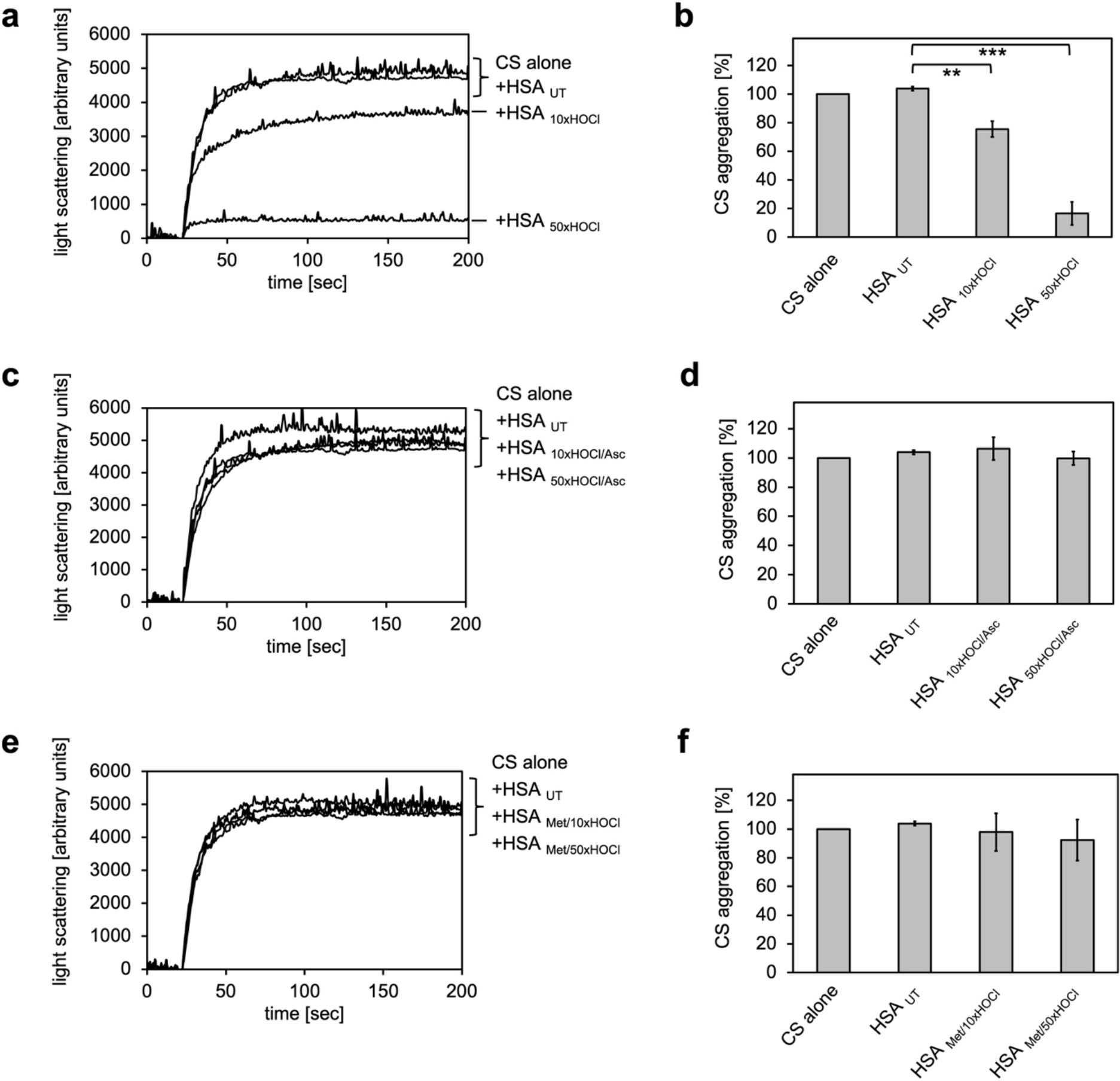
Conversion of serum albumin into a potent chaperone upon HOCl exposure is based on reversible *N*-chlorination of its basic amino acids. **(a, b)** Serum albumin, when treated with a 10- or 50-fold molar excess of HOCl (HSA_10×HOCl_ and HSA_50×HOCl_, respectively), significantly decreases aggregation of chemically-denatured citrate synthase as measured by light scattering at 360 nm. **(c, d)** Reduction of HOCl-treated HSA with a 50-fold molar excess of the antioxidant ascorbate (HSA_10×HOCl/Asc_ and HSA_50×HOCl/Asc_) switches off its chaperone activity. **(d, e)** Methylation of basic amino acid side chains prior to HOCl treatment (HSA_Met/10×HOCl_ and HSA_Met/50×HOCl_) abrogates the chaperone-like conversion of HSA. In **a**, **c** and **e** representative measurements are shown. In **b, d** and **f** data are represented as means and standard deviations from three independent experiments. Student’s t-test: **p < 0.01, ***p < 0.001. Aggregation of citrate synthase in the absence of HSA was set to 100% and all the data are presented as percentage of this control. Labels of aggregation curves are written in the order of the final intensity of light scattering of the respective treatment.

### Chaperone-like conversion of serum albumin can be reversed by antioxidants

It is well known that exposure to high HOCl concentrations, such as those present at sites of chronic inflammation, can lead to plasma protein unfolding and the formation of carbonylated and di-tyrosine cross-linked protein aggregates that cannot return to the free, functional pool of proteins upon reduction by antioxidants ^16^. Such irreversibly misfolded plasma proteins could principally act as chaperones and bind to and prevent the aggregation of other unfolding substrates through hydrophobic interactions with the newly exposed hydrophobic protein surfaces.

However, based on the observation that the chaperone activity of serum albumin increased in a dose-dependent manner with the quantity of HOCl added, we asked whether the activation of HSA chaperone function upon treatment with a 10- or 50-fold molar excess of HOCl could be mediated by reversible *N*-chlorination instead. We have found this mechanism of action in the bacterial protein RidA ^17^. We thus exposed HOCl-treated HSA to a 50-fold molar excess of the antioxidant ascorbate, which specifically removes *N*-chlorination. Indeed, ascorbate rendered HSA_10×HOCl_ and HSA_50×HOCl_ unable to prevent citrate synthase aggregation when added at the same molar excess as the HOCl-treated protein samples (Fig. 2 c, d). This result strongly supports the idea that HOCl-mediated activation of HSA chaperone function involves reversible chlorination of its side chain amines.

### Methylation of basic amino acid residues in serum albumin inhibits HOCl-induced activation of its chaperone function

Prompted by the observation that HOCl-mediated conversion of serum albumin into a potent chaperone could be reversed by ascorbate, we assumed an *N*-chlorination-based activation mechanism of its chaperone function. To further confirm this hypothesis, we blocked free amino groups of lysine and nitrogens in the guanidino-moiety of arginine residues in HSA via selective methylation. Exposure of methylated HSA to 10- or 50-fold molar excess of HOCl did not convert HSA into a chaperone, suggesting that activation of the HSA chaperone function indeed requires chlorination of its basic amino acids (Fig. 2 e, f).

### Decreased amino group content of HSA upon HOCl treatment is accompanied by an increased overall hydrophobicity of the protein

Our combined data strongly suggested that the chaperone function of HSA is activated by chlorination of its basic amino acids. HSA in its secreted form possesses 84 potential targets for chlorination by HOCl: 59 lysine residues, 24 arginine residues and one amino group at the N-terminus ^60^. To investigate the extent to which HOCl decreases the total amount of accessible, non-modified amino groups in HSA, we analyzed the free amino group content before and after HOCl-treatment using fluorescamine ^54^. Evidently, exposure to HOCl resulted in some loss of free amino groups. Amino group content of HSA decreased by approximately 10% after treatment with a 10-fold molar excess of HOCl (HSA_10×HOCl_) and by 40% upon exposure to a 50-fold molar excess of HOCl (HSA_50×HOCl_) (Fig. 3 a, b). Reduction of both chlorinated HSA samples with ascorbate resulted in a full recovery of accessible amino groups with concomitant loss of chaperone activity (Fig. 2 c, d). Activation of HSA chaperone function by HOCl thus coincides with a decrease in free amino group content, providing further evidence for an *N*-chlorination based-mechanism.

**Figure 3:**
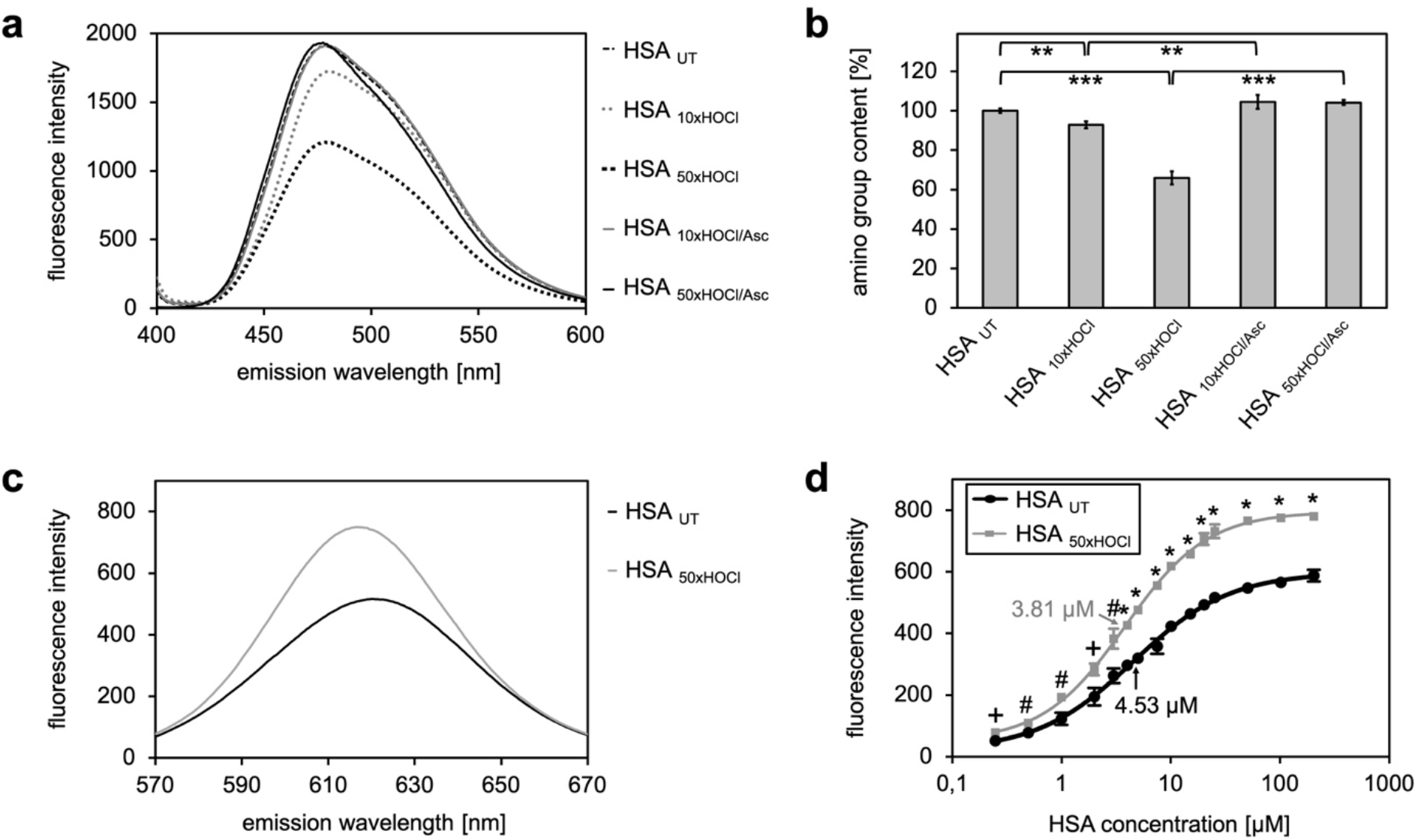
*N*-chlorination of serum albumin decreases accessible amino group content and increases surface hydrophobicity. (**a**, **b**) Amino group content of variously treated HSA was analyzed using fluorescamine. Treatment of HSA with HOCl resulted in a dose-dependent loss of free amino groups. Reduction of chlorinated HSA with ascorbate fully restored the free amino group content. (**c**) Fluorescence of 1.6 μM Nile red in the presence of native HSA or HSA, that has been treated with a 50-fold molar excess of HOCl. An increased absolute fluorescence and a shift in maximum emission wavelength (from 621.5 nm to 616.5 nm) can be observed for HSA_50×HOCl_. (**d**) Absolute fluorescence of Nile red measured at the emission maximum at 621.5 nm and 616.5 nm, respectively, was plotted against the corresponding HSA_ut_ and HSA_50×HOCl_ concentrations, respectively. The average concentrations, at which HSA_ut_ and HSA_50×HOCl_ have been half-saturated with Nile red are marked by arrows. Means and standard deviations in **b** and **d** are based on three independent experiments. (**b**) Student’s t-test: *p < 0.05, **p < 0.01, ***p < 0.001. (**d**) Student’s t-test: ^+^p < 0.05, ^#^p < 0.01, *p < 0.001. For **a** and **c** representative measurements are shown.

We argued that the reduction of positive charges on the protein’s surface through HOCl-induced *N*-chlorination should lead to an increase in surface hydrophobicity, thus allowing high-affinity binding to unfolded proteins.

To detect changes in HSA’s surface hydrophobicity upon HOCl treatment, we used the uncharged hydrophobic dye Nile red and measured its fluorescence upon addition to 25 μM native HSA or 25 μM HSA_50×HOCl_ 61. Absolute fluorescence of 1.6 μM Nile red was higher for HSA_50×HOCl_ when compared with untreated HSA with a maximum at 616.5 nm (621.5 nm for native HSA) showing a blue shift in emission maximum consistent with a decreased polarity of the protein solution (Fig. 3 c). The concentration of native HSA and HSA_50×HOCl_, at which the proteins have been half-saturated with dye were calculated (Fig. 3 d). This concentration was significantly higher for untreated HSA compared to HSA_50×HOCl_, pointing towards an increased hydrophobicity of HSA_50×HOCl_.

### Activation of neutrophil-like cells by HOCl-treated serum albumin is based on reversible *N*-chlorination

Several lines of evidence point toward a key role of HOCl-modified serum albumin in the progression of chronic inflammation, a hallmark of various degenerative diseases ^46,47^. Upon exposure to high doses of HOCl, as those present in chronically inflamed tissues, modified HSA was found to induce neutrophil NADPH oxidase activation reflected by an increased generation of reactive oxygen species and accompanied by degranulation ^47^.

Pathophysiological concentrations of HOCl are able to induce different modifications on plasma proteins including carbonylation, *N*-chlorination, cysteine and methionine oxidation or inter- and intramolecular di-tyrosine cross-linking ^8,9,11,16^, with most of them being considered irreversible. So far, it has not been elucidated which HOCl-induced modification is sufficient to convert HSA into a potent activator of leukocytes. We thus wondered whether this functional conversion of HSA might be specifically mediated by *N*-chlorination and thus reflects a reversible process, similar to the activation of its chaperone function in response to HOCl stress. To test this, we analyzed the effect of HOCl-treated HSA before and after reduction by antioxidants on the activity of the phagocytic NADPH oxidase. For this purpose, we chose the human myeloid cell line PLB-985 that acquires a neutrophil-like phenotype upon differentiation with DMSO and IFNγ^55,56^.

Generation of oxidants by the NADPH oxidase was assessed by lucigenin, a well characterized and frequently used chemiluminescence probe which predominantly reacts with superoxide anion radicals to form a light-emitting species ^62^. Differentiated PLB-985 cells, when incubated in buffer in the absence of any activating agents, showed a low basal level of superoxide generation resulting from IFNγ-mediated enhancement of the NADPH oxidase activity during the differentiation period ^63^ (Fig. 4 a). In line with expectations, treatment of the cells with phorbol-12-myristate-13-acetate (PMA), a known activator of neutrophil NADPH oxidase, led to a drastic increase in superoxide production and was thus used as positive control ^64^ (Fig. 4 a, c). Addition of untreated HSA significantly increased the generation of superoxide by > 20% compared to the mock control. No further enhancement of the NADPH oxidase activity was observed with HSA after exposure to a 10-fold molar excess of HOCl, indicating that the chosen HOCl concentration might have been too low for the conversion of HSA into a potent activator of neutrophil-like cells. In contrast, cells subjected to HSA that has been treated with a 50-fold molar excess of HOCl exhibited a substantial increase in superoxide production within the first ~ 25 minutes of incubation. In comparison to PMA, NADPH oxidase activation by HSA_50×HOCl_ proceeded at a lower rate and was less sustained, returning to basal level 40 minutes after the addition of HSA50×HOCl to the cells.

**Figure 4:**
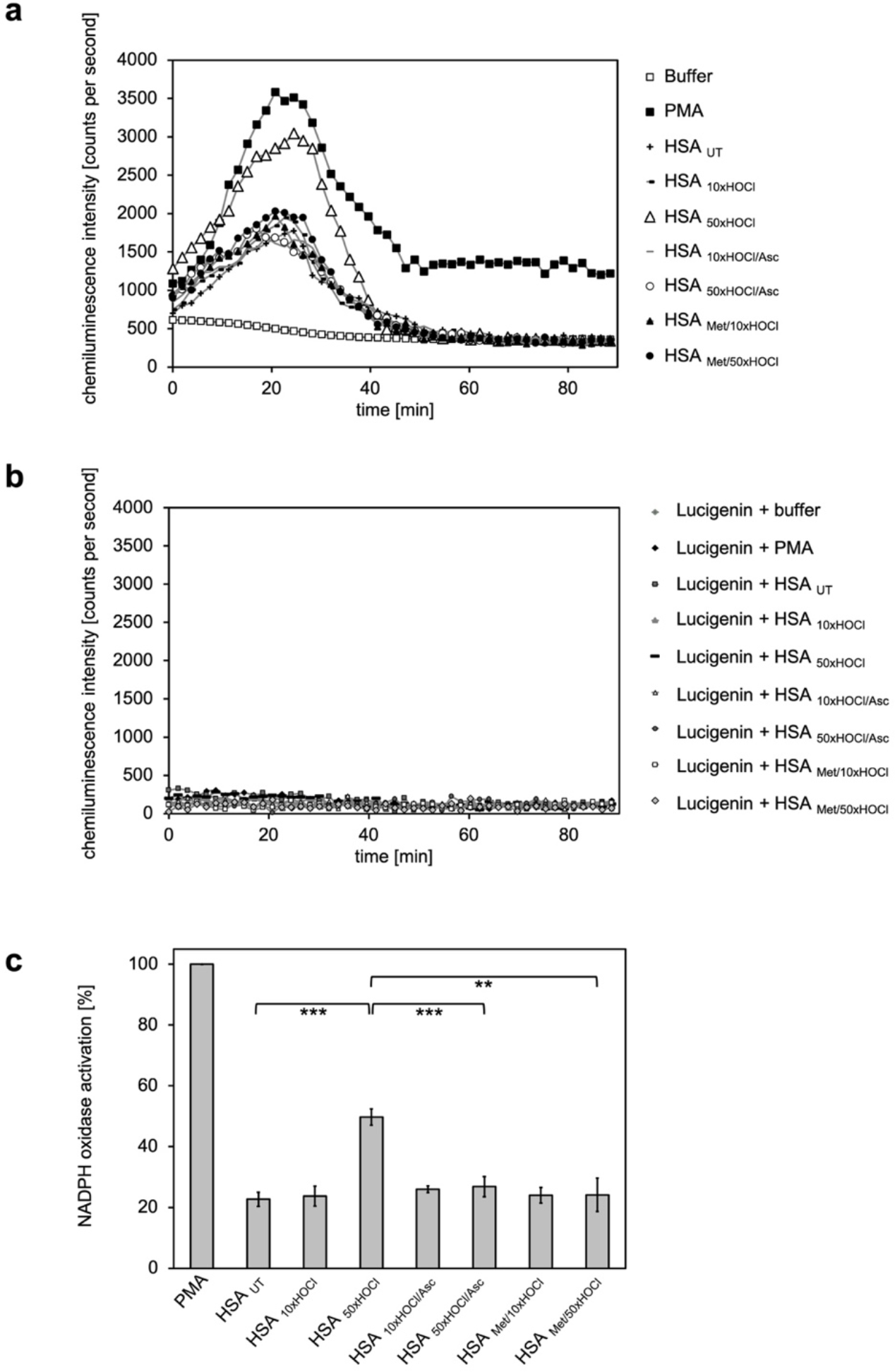
Activation of neutrophil-like cells by HOCl-treated serum albumin is mediated by reversible *N*-chlorination. Treatment with a 50-fold molar excess of HOCl (HSA _50×HOCl_) converted HSA into an efficient inducer of the neutrophil respiratory burst, reflected by the increased production and release of oxidants that induce lucigenin chemiluminescence. This activating function of HSA _50×HOCl_ could be reversed by reduction with the antioxidant ascorbate (HSA _50×HOCl/Asc_) and was abrogated by methylation of its basic amino acid side chains prior to HOCl exposure (HSA _Met/50×HOCl_). **(a)** Extracellular oxidant production by neutrophil NADPH oxidase was measured in one- to two-minutes intervals over 90 minutes at 37 °C using lucigenin-enhanced chemiluminescence. Phorbol 12-myristate 13-acetate (PMA; final concentration, 0.2 µM), untreated and variously treated HSA samples (final concentration, 3 mg ⋅ mL^−1^) from the previous citrate synthase aggregation assays (see above) or PBS buffer (basal oxidant production) were added to **(a)** differentiated PLB-985 cells in PBS buffer or **(b)** cell-free PBS puffer containing 400 µM lucigenin immediately prior to chemiluminescence measurement. **(c)** Results shown in **a** are expressed as integrated total counts (means and standard deviations of three independent measurements) higher than buffer control. Student’s t-test. **p < 0.01, ***p < 0.001. PMA-induced activation of NADPH-oxidase was set to 100%.

During our experiments we observed that *N*-chlorinated HSA can directly react with the fluorescent dye, 2’, 7’-dichlorodihydrofluorescein diacetate (H_2_DCF-DA), which is the most widely used probe for measuring intracellular ROS production ^65^, eliciting a positive signal in the absence of cells (Supplementary Fig. S1). To confirm that the observed chemiluminescent signal emitted by lucigenin derives from HSA_50×HOCl_-induced activation of the phagocytic NADPH-oxidase rather than from a similar direct reaction of lucigenin with HSA_50×HOCl_ - derived chloramines, we incubated lucigenin with the various agents in the absence of cells and no significant chemiluminescence was observed (Fig. 4 b).

Importantly, when reduced by ascorbate, HSA_50×HOCl_ lost its stimulatory effect, showing that the functional conversion of HSA to an efficient activator of neutrophil-like cells upon HOCl exposure reflects a reversible process. Since ascorbate specifically reduces *N*-chloramines without affecting other HOCl-induced modifications, these results strongly support an *N*-chlorination-based mechanism. The latter finding was further corroborated by methylation of the basic amino acids of HSA prior to HOCl treatment, which completely prevented HOCl-treated HSA from activating the neutrophil oxidative burst.

*N*-chlorinated HSA, formed upon exposure to high doses of HOCl, can be thus considered as an inflammatory response modulator that contributes to the activation of neutrophils at sites of inflammation.

### All plasma fractions tested exhibit chaperone activity upon HOCl treatment

Our finding that the major protein in human plasma, serum albumin. can be converted into a potent chaperone upon HOCl-mediated *N*-chlorination, prompted us to ask whether also other plasma protein fractions exhibit similar chaperone activity upon treatment with HOCl.

We thus tested the γ-globulin fraction, the Cohn fraction IV (comprising α- and β-globulins), and specifically α_2_-macroglobulin, an important protease inhibitor in human plasma, for chaperone activity upon exposure to various doses of HOCl.

Aside from its known interaction with proteases, native α_2_-macroglobulin (α_2_M) acts as an extracellular chaperone that binds to and prevents the accumulation of misfolded proteins, particularly during the innate immune response. Exposure to HOCl was found to further improve the chaperone function of α_2_M, but the mechanism remained unclear ^32^.

To test whether the previously observed effect is due to *N-*chlorination, we treated α_2_M with a 10- and 50-fold molar excess of HOCl (corresponding to 0.3 mM and 1.5 mM HOCl, respectively), followed by the addition of the antioxidant ascorbate for re-reduction. In line with expectations, chaperone activity of α_2_M increased with the amount of HOCl added (Fig. 5 a, b). Treatment with ascorbate, as well as methylation of basic amino acid residues prior to HOCl exposure fully inhibited the chaperone activation of HOCl-modified α_2_M. These results strongly support the notion that HOCl-induced *N*-chlorination of basic amino acid side chains also accounts for the increased chaperone activity of α_2_M.

**Figure 5:**
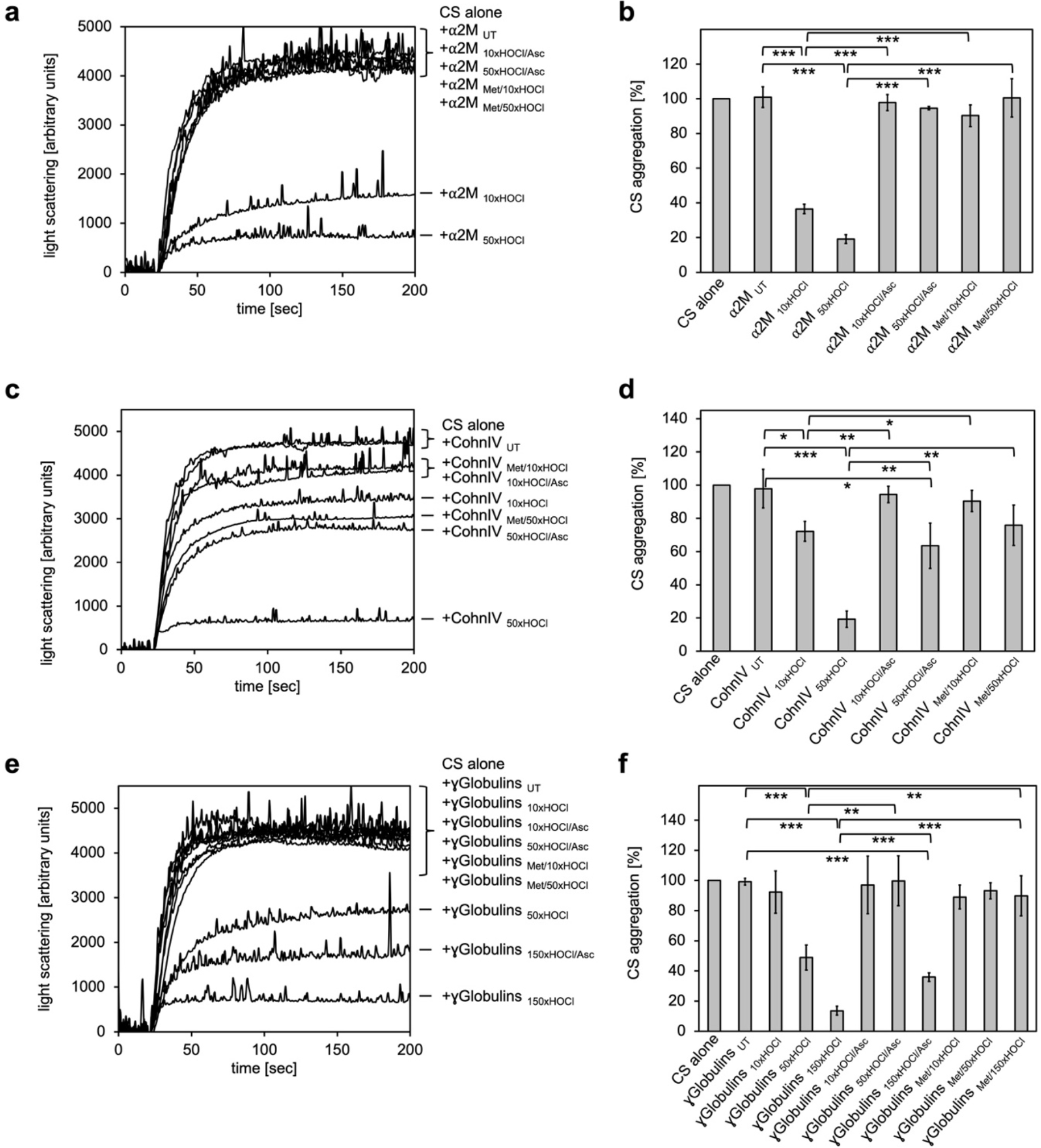
All plasma protein fractions tested exhibit reversible chaperone activity upon modification by HOCl. α_2_-Macroglobulin (**a**, **b**), Cohn fraction IV (**c**, **d**) and the γ-globulin fraction (**e**, **f**) were analyzed for chaperone activity in a citrate synthase aggregation assay upon treatment with various doses of HOCl. Each plasma protein fraction, when treated with a 10-, 50- or 150-fold molar excess of HOCl, significantly decreased aggregation of chemically-denatured citrate synthase as measured by light scattering at 360 nm. Exposure of the various HOCl-treated plasma proteins to the reductant ascorbate significantly decreased or completely inhibited their chaperone function. Methylation of basic amino acid residues prior to HOCl treatment mostly prevented chaperone-like conversion of the plasma proteins. In **a**, **c** and **e** representative measurements are shown. In **b, d** and **f** data are depicting means and standard deviations from three independent experiments. Student’s t-test: *p < 0.05, **p < 0.01, ***p < 0.001. Aggregation of citrate synthase in the absence of any plasma protein fraction was set to 100% and all the data are presented as percentage of this control. Labels of aggregation curves are written in the order of the final intensity of light scattering of the respective treatment.

Intriguingly, we also observed similar chaperone-like conversion upon HOCl treatment for Cohn fraction IV and the γ-globulin fraction (Fig. 5 c-f). In both cases, however, efficient activation of the chaperone activity required higher HOCl concentrations. It is important to note, however, that all HOCl concentrations used were below the range known to be generated directly at sites of inflammation ^31^. While exposure of Cohn fraction IV to an estimated 10-fold molar excess of HOCl (corresponding to 2 mM HOCl) had only little effect on citrate synthase aggregation, treatment with a 50-fold molar excess of HOCl strongly activated chaperone-like properties (Fig. 5 c, d). This chaperone activity was markedly reduced, but not completely inhibited, after reduction with ascorbate. Likewise, methylation of amine side chains prior to HOCl exposure did not fully abrogate the chaperone-like conversion of this protein fraction. Similarly, activation of the chaperone function of the γ-globulin fraction occurred at higher HOCl concentrations ranging from 4.3 mM to 13 mM (corresponding to an estimated 50- to 150-fold molar excess of HOCl). Strikingly, while chaperone activity increased with the amount of HOCl added, reversibility of the chaperone function by ascorbate decreased (Fig. 5 e, f).

These results suggest that at least some proteins in both plasma fractions tested are transformed to chaperones upon modification by HOCl. Efficient activation of their chaperone function is strongly, but, unlike HSA, not exclusively linked to *N*-chlorination, suggesting some other HOCl-induced modifications that cannot be removed by ascorbate in these plasma proteins.

### HOCl-induced *N*-chlorination converts the majority of plasma proteins into activators of neutrophil-like cells

Ours and others findings indicate that *N*-chlorinated serum albumin is a key factor for the stimulation of leukocytes at sites of inflammation. To test if also other plasma proteins exert this function upon HOCl exposure, we analyzed the NADPH oxidase-dependent generation of superoxide by differentiated PLB-985 cells in the presence of various treated plasma protein fractions, as described above.

Addition of α_2_-macroglobulin, treated with a 50-fold molar excess of HOCl (i.e. 1.5 mM HOCl), did not enhance superoxide production (Fig. 6 a, b). In contrast, exposure to the same molar excess of HOCl, corresponding to 10 mM HOCl in this case, converted at least some proteins of Cohn fraction IV into highly efficient stimulators of the neutrophil NADPH oxidase (Fig. 6 c, d). Analysis of the kinetics showed that activation by this HOCl-treated protein fraction occurred at a similar rate as the activation by PMA. Such a stimulatory effect was also observed for the γ-globulin fraction upon treatment with an estimated 150-fold molar excess of HOCl (i.e. 13 mM). The minimum HOCl concentration required for the activating function of these plasma proteins was thus 10 mM and conformed to the range of HOCl concentrations expected under inflammatory conditions. Importantly, the activating effect of both HOCl-treated plasma fractions was completely abolished upon reduction with ascorbate or by prior methylation of their amino groups, strongly suggesting that *N*-chlorination is the responsible mechanism for the functional switch of these proteins into efficient activators of neutrophil-like cells as well.

**Figure 6:**
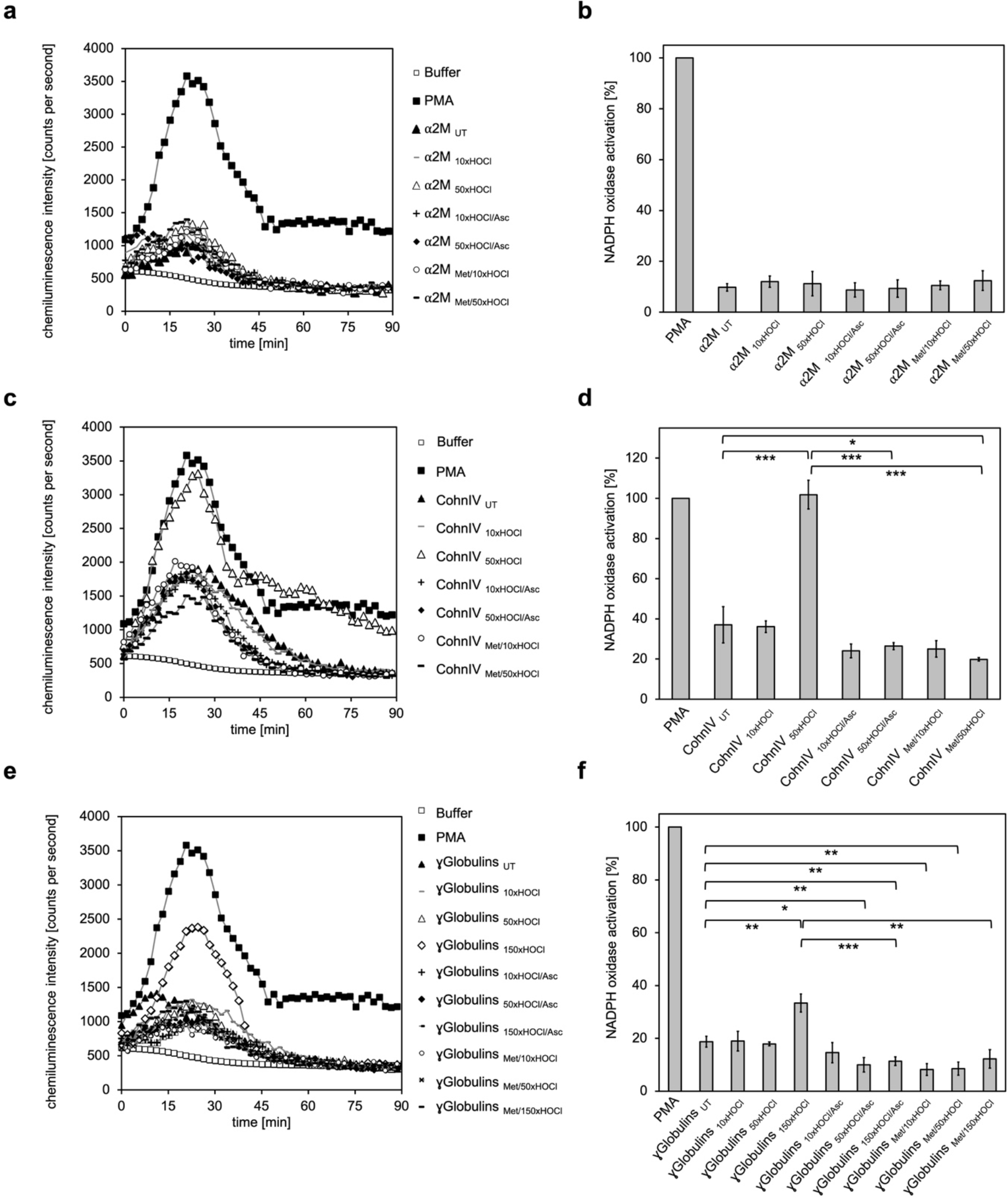
The majority of human plasma proteins stimulate neutrophil respiratory burst upon *N*-chlorination by HOCl. The effect of HOCl-treated α_2_-macroglobulin (α_2_M) (**a**, **b**), Cohn fraction IV (**c**, **d**) and the γ-globulin fraction (**e**, **f**) on the activity of the neutrophil NADPH oxidase was investigated. α_2_M, when treated with various doses of HOCl, had no influence on ROS generation by the NADPH oxidase (**a**, **b**). Treatment with a 50- or 150-fold molar excess of HOCl converted at least some proteins of Cohn fraction IV (CohnIV _50×HOCl_) and the γ-globulin fraction (γGlobulins _150×HOCl_), respectively, into efficient inducers of the neutrophil respiratory burst, reflected by the increased production and release of oxidants that induce lucigenin chemiluminescence (**c-f**). The activating function of CohnIV _50×HOCl_ and γGlobulins _150×HOCl_ could be reversed by treatment with the reductant ascorbate (CohnIV _50×HOCl/Asc_ and γGlobulins 1_50×HOCl/Asc_) and was abrogated by methylation of basic amine side chains prior to HOCl exposure (CohnIV Met/_50×HOCl_ and γGlobulins Met/1_50×HOCl_). **(a**, **c**, **e)** Extracellular oxidant production by neutrophil NADPH oxidase was measured in one- to two-minutes intervals over 90 minutes at 37 °C using lucigenin-enhanced chemiluminescence. Phorbol 12-myristate 13-acetate (PMA; final concentration (fc), 0.2 µM), untreated and the variously treated plasma fraction samples (fc, 2 mg ⋅ mL^−1^ for α_2_-macroglobulin and 3 mg ⋅ mL^−1^ for Cohn fraction IV and the γ-globulin fraction) from the previous citrate synthase aggregation assays (see above) or PBS buffer (basal oxidant production) were added to differentiated PLB-985 cells in PBS buffer containing 400 µM lucigenin immediately prior to chemiluminescence measurement. **(b**, **d**, **f)** Results are expressed as integrated total counts (means and standard deviations of three independent measurements) higher than buffer control. Student’s t-test: *p < 0.05, **p < 0.01, ***p < 0.001. PMA-induced activation of NADPH-oxidase was set to 100%.

These findings demonstrate that HOCl-mediated *N*-chlorination constitutes a key mechanism to increase the immunogenicity of plasma proteins under inflammatory conditions. Upon modification, not only serum albumin, but the majority of the plasma fractions tested, form a feed-forward inflammatory loop to amplify and sustain inflammatory responses which can lead to accelerated pathogen clearance but could also contribute to chronic inflammation.

### Activation of NADPH oxidase by *N*-chlorinated serum albumin and immunoglobulin G occurs predominantly via PI3K-dependent signaling pathways

Activity of the NADPH oxidase complex of neutrophils is regulated by several signaling pathways downstream of cell surface receptors. Central amongst these are PLC/PKC-(phospholipase C/protein kinase C) and PI3K-(phosphoinositide 3-kinase)-dependent pathways, blockade of which severely lowers NADPH oxidase activation by various stimuli such as chemotactic peptides, opsonized particles or phorbol esters ^66–69^.

To identify the predominant signaling mechanism through which the *N-*chlorinated serum albumin and the major component of the γ-globulin fraction, immunoglobulin G, activate the neutrophil respiratory burst, the effect of various inhibitors on the HSA_50×HOCl_ and IgG_150×HOCl_-induced ROS generation was tested. PMA, a direct activator of conventional PKC isoforms such as PKCα and PKCβ was used as a control for PKC-dependent activation of the NADPH oxidase ^64^.

It is worth noting, that all cell suspensions contained DMSO at a final concentration of 1%. As a consequence, NADPH oxidase activation by PMA, HSA_50×HOCl_ or IgG_150×HOCl_ proceeded at a lower rate compared to the previously described experiments (see Figs. 4 and 6), peaking at 35-50 minutes after the addition of these agents to the cells.

Pretreatment of neutrophil-like cells with 10 μM diphenyleneiodonium (DPI), a direct inhibitor of the NADPH oxidase complex, for 30 minutes prior to stimulation with PMA, HSA_50×HOCl_ or IgG_150×HOCl_ fully inhibited ROS generation confirming that the ROS production induced by these stimulatory agents was NADPH-oxidase dependent (Fig. 7). In line with expectations, presence of the PI3K inhibitor wortmannin (100 nM) had only little effect on the PMA-mediated activation of the NADPH oxidase (Fig. 7 c, d). In contrast, wortmannin strongly attenuated the HSA_50×HOCl_- and IgG1_50×HOCl_-induced ROS generation by the immune cells, suggesting a PI3K-dependent mechanism of NADPH oxidase activation (Fig. 7 a, b, d). Along this line, pretreatment with 200 nM Gö 6983, that selectively inhibits several PKC isozyme families, including the classical PKCα and PKCβII, the novel PKCα, strongly inhibited NADPH oxidase activation by PMA, but not by HSA_50×HOCl_ and IgG_150×HOCl_. These results suggest that PI3K is the key signaling component in the pathway that leads to HSA_50×HOCl_ – and IgG_150×HOCl_-dependent activation of the neutrophil NADPH oxidase.

**Figure 7:**
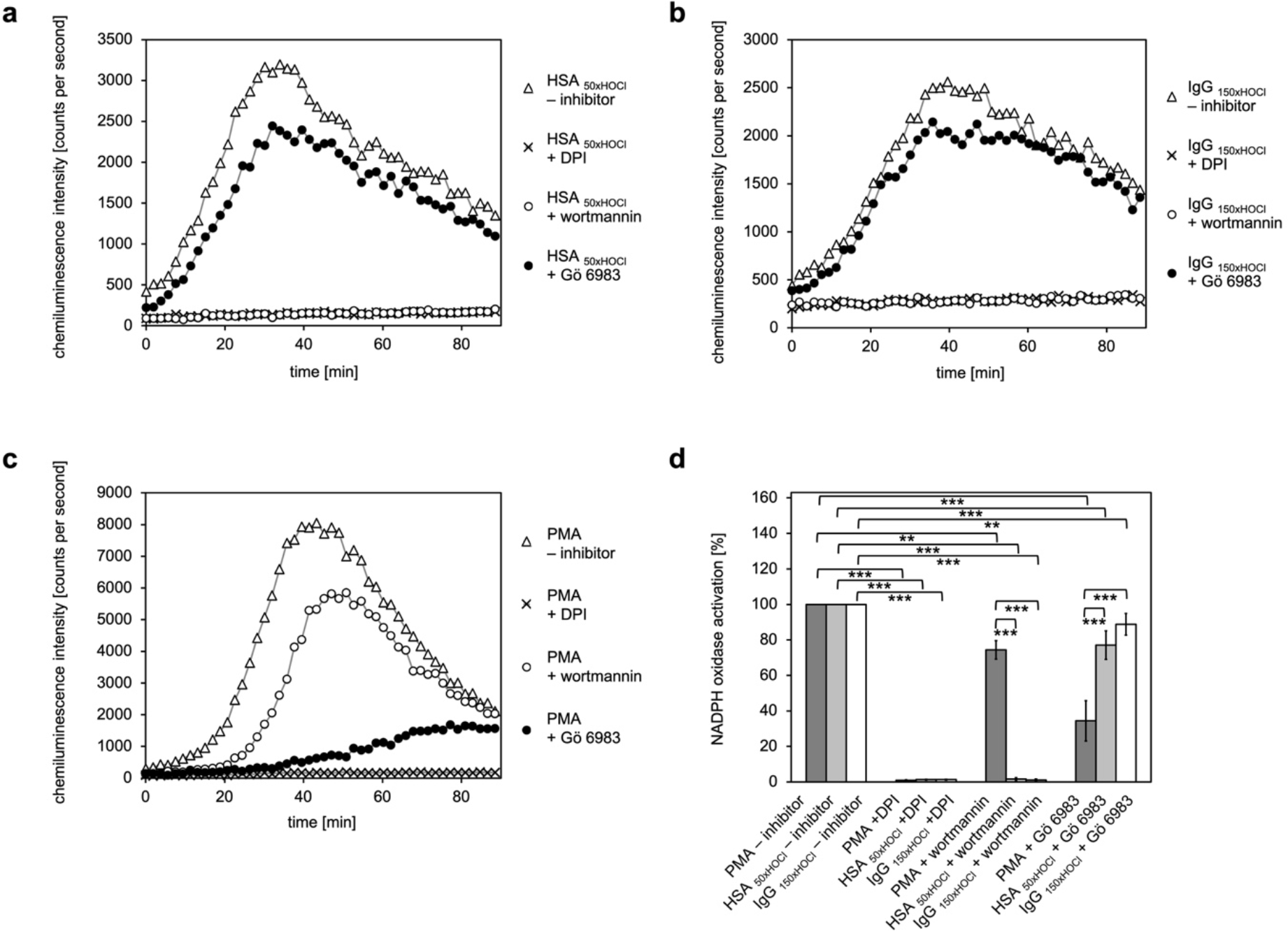
Activation of the NADPH oxidase of neutrophil-like cells by HOCl-treated serum albumin and immunoglobulin G occurs predominantly via a PI3K-dependent signaling pathway. Effect of 10 μM diphenyleneiodonium (DPI; NADPH oxidase inhibitor), 100 nM wortmannin (PI3K inhibitor) and 200 nM Gö 6983 (protein kinase C (PKC) inhibitor) on the NADPH oxidase activation mediated by **(a)** 3 mg ⋅ mL^−1^ HSA_50×HOCl_, **(b)** 3 mg ⋅ mL^−1^ IgG_150×HOCl_ and **(c)** 0.2 µM PMA was tested. **(d)** Results shown in **a**, **b** and **c** are expressed as integrated total counts (means and standard deviations of three independent measurements) higher than the respective buffer control. Student’s t-test: **p < 0.01, ***p < 0.001.

### HOCl-treated serum albumin promotes survival of neutrophil-like cells in the presence of the foreign protein antigen Ag85B, but not in the presence of staurosporine

As shown previously, activation of the NADPH oxidase by HOCl-modified serum albumin involves the action of PI3K. Since PI3K is a key component of the well-documented, anti-apoptotic PI3K/Akt signaling pathway ^70^, it was tempting to speculate that HOCl-modified HSA could promote cell survival in the presence of noxious stimuli. To test this hypothesis, we exposed neutrophil-like PLB-985 cells to the highly immunogenic mycobacterial protein antigen Ag85B and staurosporine in the presence of native or HOCl-treated HSA. The mycolyltransferase Ag85B is the major antigen produced and secreted from all mycobacterial species during infection ^71^ and has been shown to play an important role in the induction of protective immunity ^72,73^ by inducing strong T cell proliferation and IFN-γ secretion ^74,75^. Activation of neutrophils during mycobacterial infections is often accompanied by accelerated apoptosis ^76,77^, but the mechanism by which mycobacterial species or their secreted antigens induce apoptosis has not been elucidated in detail. Staurosporine, a protein kinase inhibitor, has been characterized as an efficient inducer of apoptosis in various cell types via caspase-dependent and -independent pathways ^78,79^ and thus, was also used as an cell death-promoting agent in our experiments.

After one hour or six hours of incubation with Ag85B and staurosporine, respectively, viability of the variously treated cells was evaluated by flow cytometry using Annexin V-FITC/propidium iodide (PI) staining. During early apoptosis, phosphatidylserine is translocated from the inner to the outer cell membrane leaflet with the plasma membrane left intact and thus, available for binding of extrinsically applied annexin V protein ^80^. In late apoptosis/necrosis, the integrity of the plasma membrane is lost, allowing the normally membrane-impermeable propidium iodide to enter and stain the DNA. Cells that are in late apoptosis or necrotic are thus both Annexin V-FITC and PI positive. Accordingly, cells that are viable are both Annexin V-FITC and PI negative.

Data plots were generated from analysis of ungated data (Fig. 8). Viable cells appear in the lower left quadrant (Q4), early apoptotic cells in lower right quadrant (Q3) and late apoptotic/necrotic cells in the upper right (Q2) quadrant.

**Figure 8:**
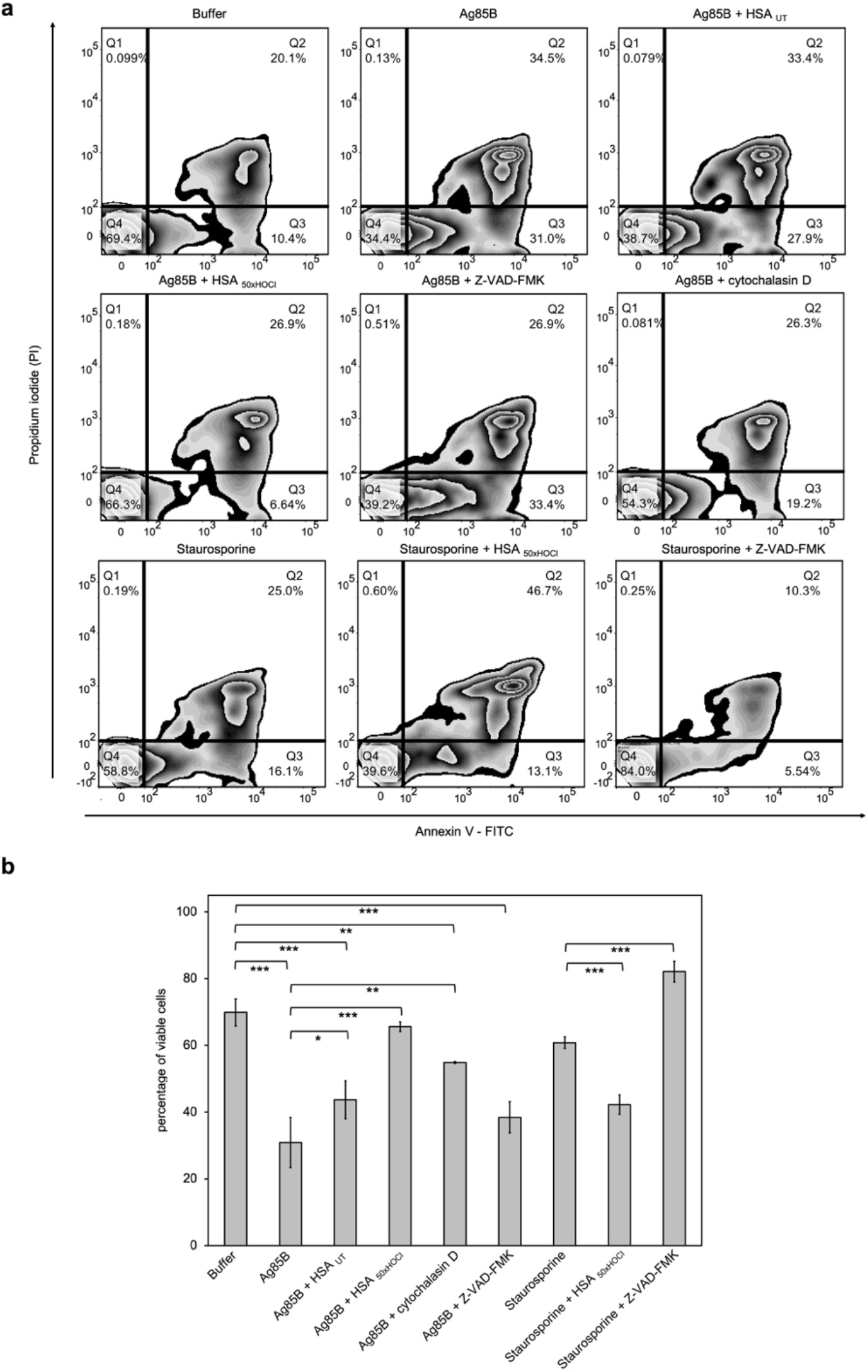
HOCl-treated serum albumin improves survival of neutrophil-like cells in the presence of the major mycobacterial protein antigen Ag85B. Differentiated neutrophil-like PLB-985 cells were preincubated with 50 μM Z-VAD-FMK, 155 μM native (HSA_UT_) or HOCl-treated HSA (HSA_50×HOCl_) prior to the addition of 1 μM Ag85B or 2 μM staurosporine. After one hour (Ag85B) or six hours (staurosporine) of incubation, viability of the variously treated cells was assessed by flow cytometry using Annexin V/propidium iodide (PI) staining. Cells treated only with buffer served as control. **(a)** Annexin V-FITC vs. propidium iodide dot plots show all analyzed events. Staining of cells simultaneously with Annexin V-FITC (green fluorescence) and the non-vital dye PI (red fluorescence) allows the discrimination of viable, intact cells (FITC-, PI-; Q4), early apoptotic (FITC+, PI-; Q3) and late apoptotic/necrotic cells (FITC+, PI+; Q2). 20,000 events were acquired and recorded per sample. Data were analyzed using FlowJo (version 10) software. Results shown are representative of three experiments. **(b)** Results of three independent experiments for viable cells are shown (means and standard deviations). Student’s t-test: *p < 0.05, **p < 0.01, ***p < 0.001.

Upon treatment with Ag85B, the ratio of viable cells to early/late apoptotic cells has markedly decreased compared to the control cells, that were incubated in the absence of Ag85B, pointing towards a lifetime-limiting effect of Ag85B. Pretreatment with the broad-spectrum pan-caspase inhibitor Z-VAD-FMK did not significantly affect Ag85B-induced cell death, suggesting that Ag85B exerts its lethal effect through a caspase-independent mechanism. To test whether the toxicity of Ag85B depends on its uptake by the immune cells, we pretreated the cells with cytochalasin D, an inhibitor of phagocytosis, prior to the addition of Ag85B. Reduction of the cells’ phagocytic capacity resulted in markedly improved cell survival in the presence of Ag85B. This phenomenon is illustrated by a shift of the cells from Q3 to Q4 on the Annexin V-FITC/PI plot with concomitant reduction of necrotic cells. Remarkably, pretreatment with HOCl-modified HSA, added at a 155-fold molar excess over Ag85B, completely prevented cell death upon addition of Ag85B. In contrast, native HSA had almost no effect on Ag85B-induced cell death. We thus speculated that HOCl-treated HSA could rescue immune cells from Ag85B-induced cell death by preventing or strongly reducing its uptake, rather than by boosting PI3K/Akt signaling. In support of this conclusion, Z-VAD-FMK, but not HOCl-treated HSA was able to significantly reduce staurosporine-induced apoptosis. Instead, when combined with staurosporine, HOCl-modified HSA triggered necrosis with the majority of the cells being Annexin V and PI positive.

### HOCl-treated serum albumin effectively binds Ag85B and reduces its uptake by neutrophil-like cells

Prompted by the finding that HOCl-treated HSA can act as a chaperone being highly effective at preventing protein aggregation, we asked whether *N*-chlorinated HSA can also bind to Ag85B and thus prevent its uptake by immune cells. To test this, Ag85B was diluted stepwise in the presence of native or HOCl-treated HSA and aggregation of Ag85B was monitored by light scattering. When denatured Ag85B was diluted into buffer, it readily formed aggregates (Fig. 9 a, b). This aggregation could not be prevented by untreated HSA. Presence of HOCl-modified HSA, however, significantly reduced the aggregation of Ag85B when added at a 80-fold molar excess over the protein. Again, reduction with ascorbate fully abrogated HSA_HOCl_’s ability to bind Ag85B.

**Figure 9:**
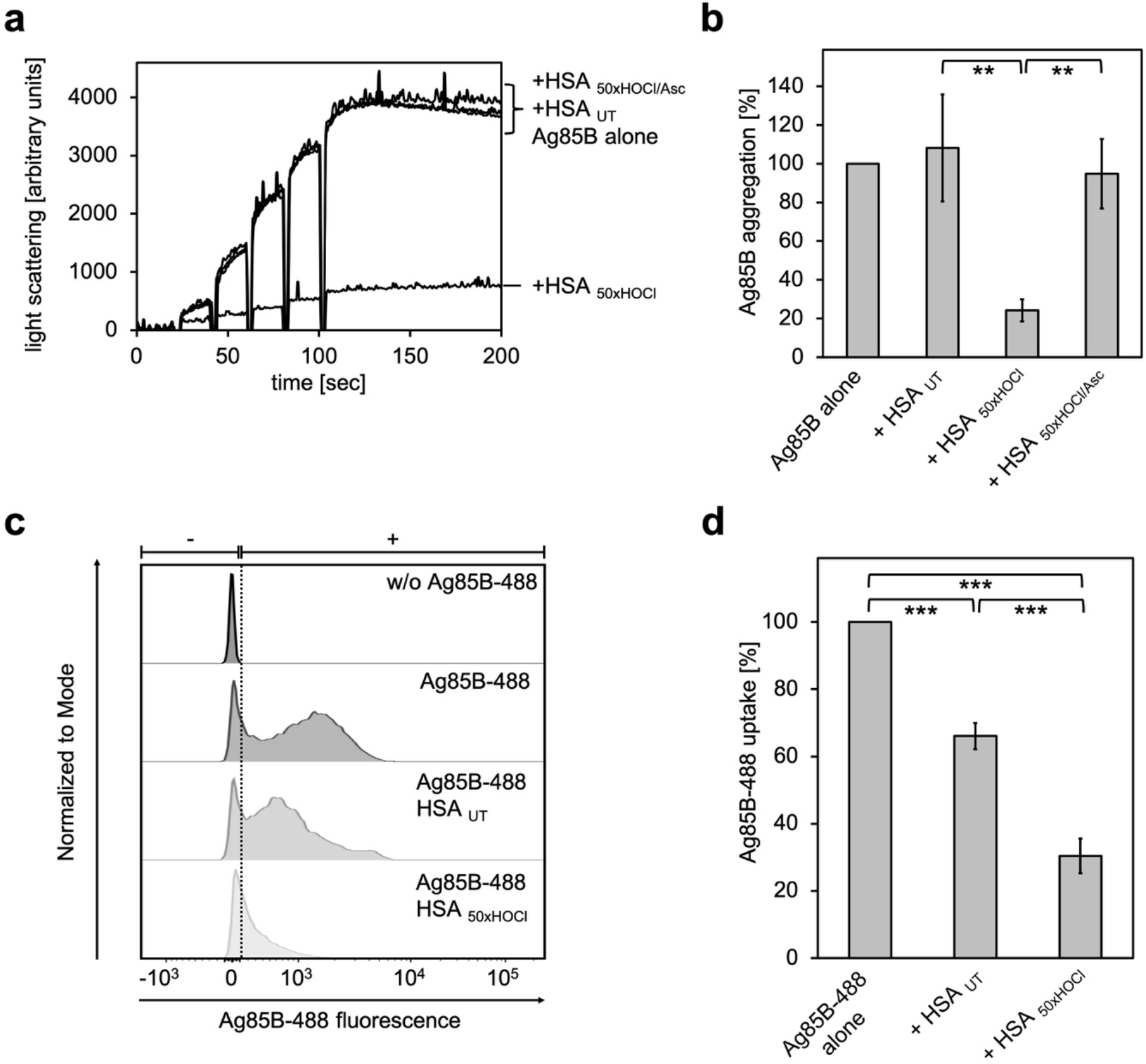
HOCl-treated serum albumin binds to and prevents uptake of the major mycobacterial protein antigen Ag85B by neutrophil-like cells. **(a, b)** HSA, treated with a 50-fold molar excess of HOCl (HSA_50×HOCl_) significantly decreased aggregation of denatured Ag85B as measured by light scattering at 360 nm. Reduction of HSA_50×HOCl_ with a 50-fold molar excess of the antioxidant ascorbate (HSA_50×HOCl/Asc_) reversed this chaperone activity. **(a)** A representative measurement of Ag85B aggregation in the presence of native HSA (HSA_UT_), HSA_50×HOCl_ and HSA_50×HOCl/Asc_ is shown. Labels of aggregation curves are written in the order of the final intensity of light scattering of the respective treatment. **(b)** Data are represented as means and standard deviations from three independent aggregation assays. Aggregation of Ag85B in the absence of HSA was set to 100% and all the data are presented as percentage of this control. **(c, d)** Differentiated neutrophil-like PLB-985 cells were incubated in the absence or presence of the fluorescently-labeled Ag85B protein, Ag85B-488. In some cases, HSA_UT_ or HSA_50×HOCl_ was added at a 50-fold molar excess over Ag85B-488 to the cells. After one hour of incubation, uptake of fluorescent Ag85B-488 by the variously treated cells was assessed by flow cytometry. 30,000 events were acquired and recorded per sample. Data were analyzed using FlowJo (version 10) software. **(c)** Single-parameter histogram overlays of Ag85B-488 fluorescence of the various samples are shown. **(d)** Results of three independent experiments are shown (means and standard deviations). Student’s t-test: **p < 0.01, ***p < 0.001. Uptake of Ag85B-488, reflected by the median fluorescence intensity, in the absence of HSA was set to 100%.

We argued that Ag85B’s association with HOCl-modified HSA could decrease its propensity to enter immune cells and thus, provide a possible explanation for the enhanced survival of neutrophils in the presence of this mycobacterial antigen. To test whether this HSA_HOCl_ - mediated cell survival is indeed due to a reduced uptake, we added recombinant fluorescently-labeled Ag85B (Ag85B-488) to differentiated neutrophil-like cells and analyzed its uptake in the presence of native or HOCl-treated HSA using flow cytometry. Cells conincubated with native HSA accumulated around 30% less Ag85B compared to cells treated with Ag85B alone (Fig. 9 c, d). But presence of HOCl-treated HSA reduced uptake of Ag85B by up to 80%. Sustained cell viability in the presence of Ag85B is thus most likely linked to decreased antigen uptake by the immune cells.

## DISCUSSION

Myeloperoxidase (MPO)-derived HOCl is one of the strongest oxidants produced and released by neutrophils during infection and inflammation (recently reviewed in ref. ^42^). Because of its high reactivity with a wide variety of biomolecules, HOCl forms an efficient weapon against a wide range of pathogens, but also contributes to host tissue injury associated with diverse inflammatory diseases ^81^. Due to their high abundance in blood and interstitial fluid, plasma proteins constitute major targets for HOCl-mediated modification with thiol oxidation and chlorination of amine side chains being the favored reactions ^22^, followed by some irreversible modifications including tryptophan oxidation, peptide bond cleavage or di-tyrosine cross-linking ^82^, when exposed to high doses of HOCl as those present at sites of chronic inflammation. Formation and accumulation of such advanced oxidation protein products (AOPPs) is indeed a hallmark of various inflammatory diseases ^44,83^.

Aside from their role as important scavengers of the majority of generated HOCl ^39,84^, there is a growing body of evidence demonstrating protective and/or immunomodulatory function of these AOPPs during inflammation. α_2_-Macroglobulin (α_2_M), a plasma glycoprotein, already acts as an extracellular chaperone capable of suppressing the aggregation of a range of proteins ^85^, but modification by HOCl has been reported to drastically increase its chaperone activity by a yet unknown mechanism ^32^. Previously, we observed a similar phenomenon in *E. coli*, where the protein RidA, a member of the YjgF/YER057c/UK114 protein family, was converted into an efficient chaperone holdase upon exposure to HOCl by a mechanism that involved the reversible chlorination of its basic amino acids ^17^. This finding was rather surprising, since activation of other known redox-regulated chaperones, such as Hsp33 or 2-cys peroxiredoxins ^19,86^, by HOCl occured via oxidation of their cysteine residues and not by *N*-chlorination, a modification that before our study has been thought to have a mostly deleterious effect on protein function. Since similar observations were made more recently for *E. coli* CnoX (YbbN) ^25^ and the HSA homologue bovine serum albumin (BSA) ^17^, HOCl-mediated *N*-chlorination can, therefore, be considered a novel, cysteine-independent and reversible mechanism of chaperone activation in response to HOCl-stress.

Encouraged by these findings, we hypothesized that HOCl-mediated improvement of α_2_M’s chaperone function may be also based on *N*-chlorination and suspected that α_2_M could be not the only plasma protein exhibiting chaperone activity upon modification by HOCl. In fact, we found that not only α_2_M but a number of proteins in all plasma fractions tested turned into chaperone-like proteins, being highly effective at preventing formation of potentially toxic protein aggregates *in vitro*. The role of plasma proteins in inflamed tissue thus goes beyond that of a passive sink for HOCl.

Since exposure to high HOCl concentrations can lead to oxidative, irreversible protein unfolding ^16^, one might speculate that the observed chaperone activity of such unfolded plasma proteins could simply be the result of an increased affinity to other unfolded proteins. Here, we provide direct evidence that HOCl-mediated conversion of plasma proteins into potent chaperones depends primarily on reversible chlorination of their basic amino acids. Along this line, treatment of HSA with HOCl led to dose-dependent reduction of the accessible amino group content accompanied by an increase in surface hydrophobicity, providing an obvious explanation for the increased affinity to unfolding proteins of *N*-chlorinated plasma proteins. Since *N*-chlorination does not irreversibly alter the structural and functional properties of a protein, this reversible post-translational modification provides a new strategy to recruit new chaperone-like proteins on demand in response to HOCl-stress in order to minimize self-damage associated with the formation of protein aggregates during inflammation.

In recent years, other roles for HOCl-modified plasma proteins and lipoproteins in inflammatory processes have also been described. For example, HOCl-oxidized low-density lipoproteins ^87^ as well as HOCl-modified HSA ^47,48^ have been shown to elicit various polymorphonuclear leukocyte (PMNL) responses such as the NADPH-dependent generation of reactive oxygen species (ROS), degranulation or shape change ^47^, but the mechanism for the HOCl-mediated functional conversion into a potent activator of human PMNLs remained unclear.

As *N*-chlorination was the principal chemical modification responsible for the plasma protein’s switch to a chaperone-like holdase, we hypothesized that it could also be the reason for the activation of immune cells. To mimic an *in vivo* situation, where plasma proteins are located in immediate vicinity to accumulated neutrophils in inflamed tissue, we exposed several plasma fractions to HOCl at pathophysiological concentrations ^31^ and investigated the effect of the resulting products on ROS production by neutrophil-like cells. Remarkably, we found that not only the main plasma protein HSA, but the majority of plasma fractions tested, when treated with a sufficient amount of HOCl, were able to elicit a significant immune response, as shown by the increased generation of ROS by the immune cells. This effect was not seen upon re-reduction of the HOCl-modified proteins or prior methylation of their basic amino acids, strongly supporting a *N*-chlorination based mechanism as well. To the best of our knowledge, this is the first study which demonstrates that reversible HOCl-mediated *N*-chlorination is the principal mechanism of turning plasma proteins into critical modulators of the innate immune response.

Gorudko et al. reported that the phosphoinositide 3-kinase (PI3K) inhibitor wortmannin inhibited the stimulating effect of HOCl-modified HSA on immune cells ^47^. The same was true in our model system: the immunomodulatory action of both HOCl-modified HSA and immunoglobulin G could be fully inhibited or strongly attenuated by wortmannin. PI3K and its downstream effectors, such as the serine/threonine kinase Akt, are indeed considered to play a key role in the regulation of the neutrophil NADPH oxidase ^88,89^. Since the PKC inhibitor Gö 6983 showed only little effect on HSA_50×HOCl_- and IgG_150×HOCl_-mediated NADPH oxidase activation, we thus propose that these proteins stimulate the neutrophil respiratory burst predominantly via PI3K-dependent signaling pathways. However, further studies are needed to investigate the exact mechanism by which these proteins trigger PI3K signaling. A possible scenario might be, that *N*-chlorination confers higher affinity to a membrane scavenger receptor allowing the binding of HOCl-modified HSA. In support of this theory, it has been reported that *N*-chlorinated, but not native HSA can irreversibly bind to and block the major high-density lipoprotein receptor, scavenger receptor class B, type 1 (SR-BI) ^90^. It is therefore tempting to speculate that *N*-chlorination could also increase the affinity of IgG to Fc gamma receptors (FcγRs).

The PI3K/Akt pathway is also considered an anti-apoptotic pathway ^91,92^. Thus, it was tempting to hypothesize that HOCl-modified HSA may play a role as a pro-survival molecule during inflammation. Remarkably, we found that HOCl-modified HSA indeed enhances survival of neutrophils in the presence of the highly immunogenic mycobacterial protein antigen Ag85B, however, not in the presence of staurosporine, a broad spectrum protein kinase inhibitor that induces apoptosis in various cell types ^78,79^. Looking for a mechanism by which HOCl-modified HSA is able to rescue immune cells from Ag85B-induced cell death, we found that it can effectively bind to and strongly decrease the phagocytosis of Ag85B. Internalization of Ag85B by the cells proved to be the direct cause of cell death and could also be prevented by other inhibitors of phagocytosis.

Neutrophils are typically short-lived, but their apoptosis can be delayed both by microbial products and by various proinflammatory stimuli ^93,94^. In this study, we describe HOCl-modified HSA as a novel pro-inflammatory mediator, which can promote cell survival by binding to highly immunogenic foreign antigens and reducing their phagocytosis at sites of bacterial infection. A similar phenomenon was observed in other studies, where HOCl-modified HSA was shown to bind and neutralize proteins from HIV and West Nile virus ^95,96^. The immunomodulatory effects of *N-*chlorinated plasma proteins found in this and previous studies constitute a double-edged sword. Although stimulation of neutrophil respiratory burst and enhanced neutrophil survival may be beneficial for pathogen elimination at the initial stage of infection, it can eventually perpetuate a positive feedback loop and contribute to the development and progression of chronic inflammation (Fig. 10). Secretion of HOCl and other oxidants by permanently activated neutrophils leads not only to the destruction of neighboring, healthy cells resulting in tissue injury ^26,97,98^, but generates more *N*-chlorinated plasma proteins. Similarly, the pro-survival effect of *N*-chlorinated HSA is not only positive, as cell death and the subsequent recognition of dying neutrophils by macrophages has a critical function in the resolution of the inflammatory response and is strictly required to protect the surrounding tissue and prevent pathological sequelae ^99^. Were all these properties of AOPPs dependent on the numerous irreversible modifications reportedly caused by HOCl exposure, this could lead to a spreading out-of-control immune reaction. The presence of high concentrations of antioxidants such as ascorbate and glutathione in plasma, both of which can reduce *N*-chlorination, provide a mechanism to contain the immune reaction at the site of inflammation. Indeed, depletion of these antioxidants is often associated with chronic inflammation and other diseases ^100–103^ and antioxidant therapy such as high-dose intravenous vitamin C treatment leads to decrease of inflammation markers such as CRP ^104–106^. Our data explains important aspects of these effects and further highlights the role of antioxidant homeostasis in inflammatory processes.

**Figure 10:**
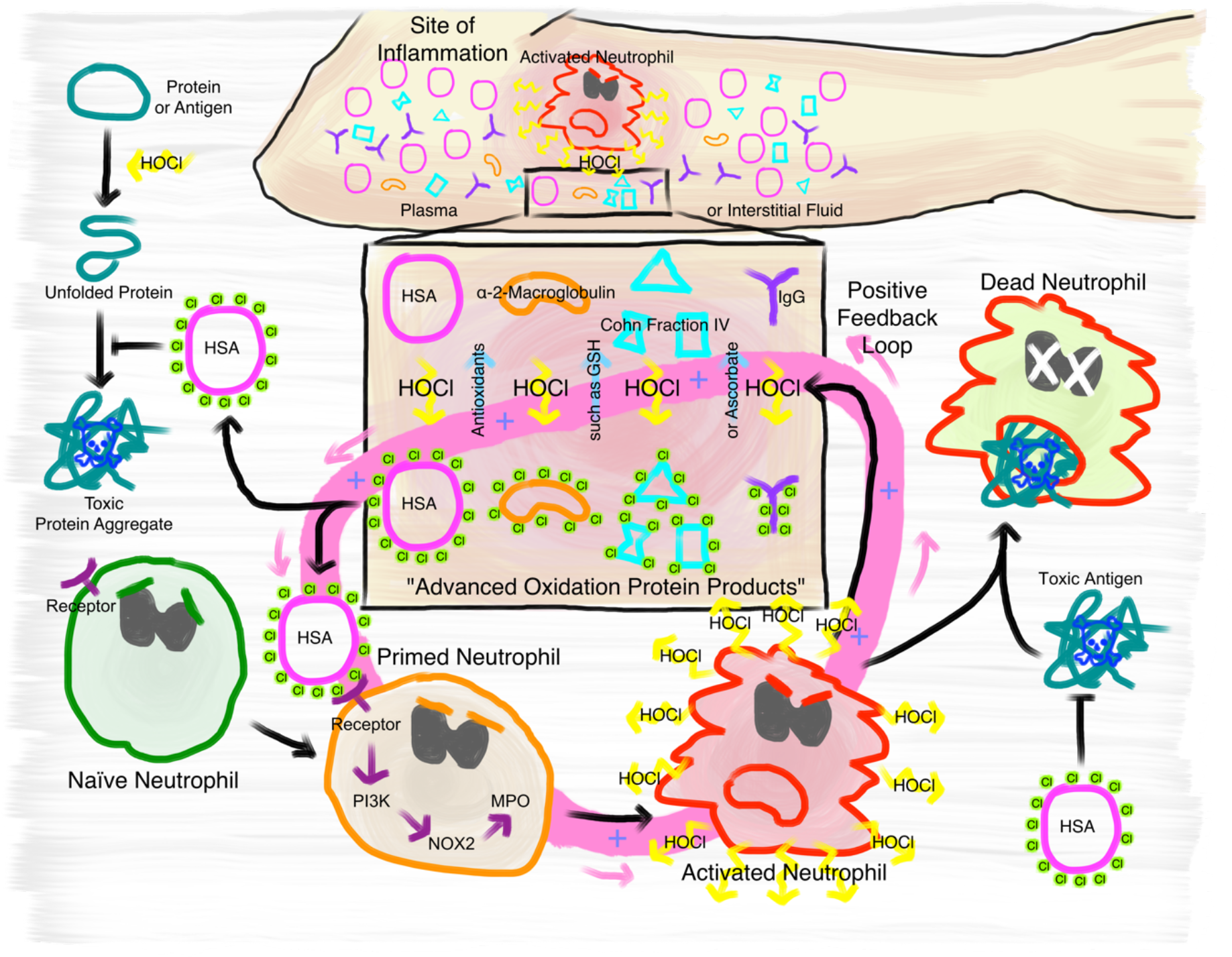
Proposed mechanism of the immunomodulatory role of *N*-chlorination of plasma proteins at a site of inflammation. At the site of inflammation neutrophils (and potentially other immune cells) are activated. Neutrophils then produce HOCl at concentrations of up to 25 to 50 mM per hour. Plasma proteins, such as HSA, α- and β-globulins (Cohn Fraction IV), γ-globulins (IgG), and α_2_-Macroglobulin then act as an effective sink. Reversible *N*-chlorination of these proteins turns them into effective chaperons, which can prevent the formation of protein aggregates and their uptake by immune cells, enhancing the survival of neutrophils in the presence of toxic antigens. *N*-chlorinated plasma proteins also activate more immune cells, which then in turn produce more HOCl, leading to the formation of more *N*-chlorinated plasma proteins. This positive feedback loop can be attenuated and deactivated by antioxidants present in plasma, such as ascorbate and reduced glutathione (GSH).

In summary, our data support a critical role for HOCl-mediated *N*-chlorination of plasma proteins during inflammatory processes and suggest that it is the critical modification mediating the physiological effects of so-called AOPPs. Although the reversible conversion of HOCl-modified plasma proteins to effective chaperones confers protection against HOCl-induced protein aggregation, the increase of their immunogenicity can potentially exacerbate self-damage at sites of inflammation through a positive feedback loop. The fact that the activation of immune cells is mediated through *N*-chlorination, a modification that is reversible by antioxidants present in plasma, provides a mechanism to attenuate or deactivate this positive feedback loop. These findings contribute importantly to our understanding of the development and progression of chronic inflammation.

## ACKNOWLEDGMENTS

Funding for this study was provided by the German Research Foundation (DFG) through grant LE2905/1-2 to LIL as part of the priority program 1710 ‘Dynamics of Thiol-based Redox Switches in Cellular Physiology’. Parts of this manuscript were written during a Writing Retreat funded by the Ruhr-Universität Bochum Research School RURS^plus^.

## CONFLICT OF INTEREST STATEMENT

The authors have no conflicting financial interests.

## AUTHOR CONTRIBUTIONS

AU, AVS, AM, and LIL conceptualized the study. AU, AVS and AM performed the experiments. AU and AVS analyzed the results. NL purified α_2_-Macroglobulin and Ag85B. AVS consulted on the manuscript and AU and LIL wrote the manuscript.

## SUPPLEMENTAL INFORMATION

**Supplementary Data:** Supplementary Figure 1 and Supplementary Materials and Methods

**Figure 1 – Source Data 1:** Numerical light scattering data obtained during protein aggregation assays represented in figure 1 a and b

**Figure 2 – Source Data 1:** Numerical light scattering data obtained during protein aggregation assays represented in figure 2 a and b

**Figure 2 – Source Data 2:** Numerical light scattering data obtained during protein aggregation assays represented in figure 2 c and d

**Figure 2 – Source Data 3:** Numerical light scattering data obtained during protein aggregation assays represented in figure 2 e and f

**Figure 3 – Source Data 1:** Numerical fluorescence spectroscopy data obtained during determination of free amino groups represented in figure 3 a and b

**Figure 3 – Source Data 2:** Numerical fluorescence spectroscopy data obtained during determination of protein hydrophobicity represented in figure 3 c

**Figure 3 – Source Data 3:** Numerical fluorescence spectroscopy intensity data obtained during determination of protein hydrophobicity represented in figure 3 d

**Figure 4 – Source Data 1:** Numerical chemiluminescence plate reader data represented in figure 4 a, b, and c

**Figure 5 – Source Data 1:** Numerical light scattering data obtained during protein aggregation assays represented in figure 5 a and b

**Figure 5 – Source Data 2:** Numerical light scattering data obtained during protein aggregation assays represented in figure 5 c and d

**Figure 5 – Source Data 3:** Numerical light scattering data obtained during protein aggregation assays represented in figure 5 e and f

**Figure 6 – Source Data 1:** Numerical chemiluminescence plate reader data represented in figure 6 a and b

**Figure 6 – Source Data 2:** Numerical chemiluminescence plate reader data represented in figure 6 c and d

**Figure 6 – Source Data 3:** Numerical chemiluminescence plate reader data represented in figure 6 e and f

**Figure 7 – Source Data 1:** Numerical chemiluminescence plate reader data represented in figure 7 a and d

**Figure 7 – Source Data 2:** Numerical chemiluminescence plate reader data represented in figure 7 b and d

**Figure 7 – Source Data 3:** Numerical chemiluminescence plate reader data represented in figure 7 c and d

**Figure 8 – Source Data 1:** Numerical flow cytometry data represented in figure 8 a and b

**Figure 9 – Source Data 1:** Numerical light scattering data obtained during protein aggregation assays represented in figure 9 a and b

**Figure 9 – Source Data 2:** Numerical flow cytometry data obtained during protein aggregation assays represented in figure 9 c and d

