## Supplementary Data for "*N*-chlorination mediates protective and immunomodulatory effects of oxidized human plasma proteins"

Supplementary data contents:

- Supplementary Material and Methods
- Supplementary Figure

### SUPPLEMENTARY MATERIAL AND METHODS

#### Detection of reactive oxygen species using the fluorescence probe $H_2$DCFDA

Intracellular production of reactive oxygen species can be monitored by the oxidation of non-fluorescent 2',7'-dichlorodihydrofluorescein diacetate ($H_2$DCF-DA) to the fluorescent 2', 7'-dichlorofluorescein (DCF). $H_2$DCF-DA (Thermo Fisher Scientific, Waltham, MA) at a final concentration of 7 µM was preincubated with 1xPBS buffer in a non-transparent, black, clear-bottom 96-well plate (Nunc, Rochester, NY) for 15 minutes at 37 °C. Fluorescence intensity was recorded every 2 minutes using the Synergy H1 multi-detection microplate reader (Biotek) at an excitation wavelength of 488 nm and an emission wavelength of 525 nm. Then 34 µM native or HOCl-treated serum albumin, or buffer were added to the wells. Fluorescence intensity was then recorded for 45 minutes.

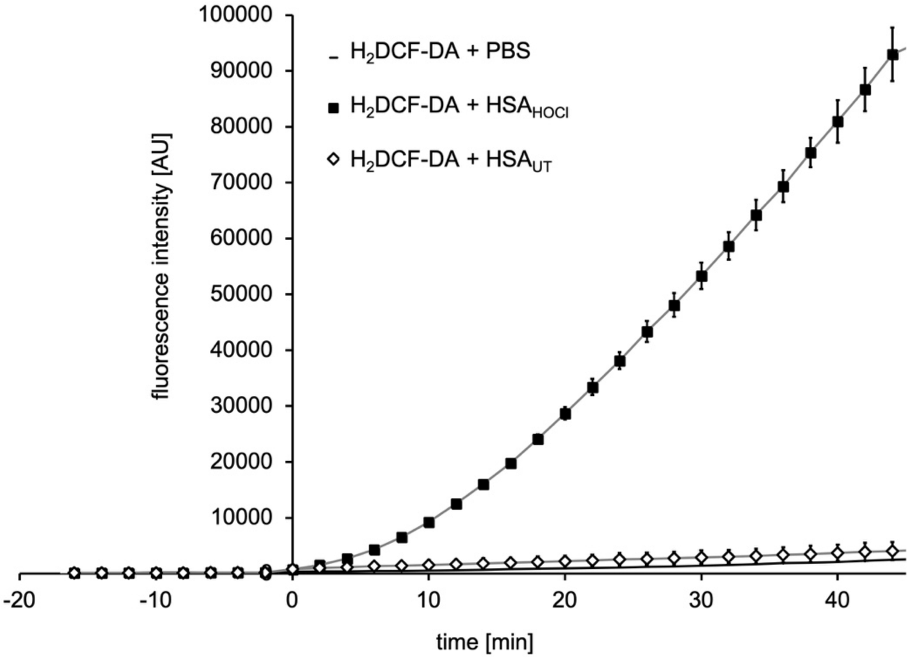


**Supplementary Figure S1: *N*-chlorinated serum albumin converts 7'-dichlorodihydrofluorescein diacetate ($H_2$DCF-DA) to the fluorescent 2', 7'-dichlorofluorescein (DCF).** $H_2$DCF-DA is a commonly used fluorescent probe for monitoring intracellular production of reactive oxygen species. $H_2$DCF-DA was preincubated with 1xPBS for 15 minutes prior to the addition of native HSA ($HSA_{UT}$), HOCl-modified HSA ($HSA_{HOCl}$) or buffer (PBS). The fluorescence intensity of DCF was recorded for 45 minutes.
